# An unconventional mechanism of IL-1β secretion that requires Type I IFN in lupus monocytes

**DOI:** 10.1101/2023.08.03.551696

**Authors:** Simone Caielli, Preetha Balasubramanian, Juan Rodriguez-Alcazar, Uthra Balaji, Zurong Wan, Jeanine Baisch, Cynthia Smitherman, Lynnette Walters, Paola Sparagana, Djamel Nehar-Belaid, Radu Marches, Lorien Nassi, Katie Stewart, Julie Fuller, Jacques F. Banchereau, Jinghua Gu, Tracey Wright, Virginia Pascual

**Author notes:** Contribute equally (Preetha Balasubramanian, Juan Rodriguez-Alcazar).

## Abstract

Systemic Lupus Erythematosus (SLE) is characterized by autoreactive B cell activation, upregulation of Type I Interferon (IFN) and widespread inflammation. Mitochondrial nucleic acids (NAs) are increasingly recognized as triggers of IFN^1^. Thus, defective removal of mitochondria from mature red blood cells (Mito^+^ RBCs), a feature of SLE, contributes to IFN production by myeloid cells^2^. Here we identify blood monocytes (Mo) that have internalized RBCs and co-express IFN-stimulated genes (ISGs) and interleukin-1β (IL-1β) in SLE patients with active disease. We show that ISG expression requires the interaction between Mito^+^ RBC-derived mitochondrial DNA (mtDNA) and cGAS, while IL-1β production entails Mito^+^ RBC-derived mitochondrial RNA (mtRNA) triggering of RIG-I-like receptors (RLRs). This leads to the cytosolic release of Mo-derived mtDNA that activates the NLRP3 inflammasome. Importantly, IL-1β release depends on the IFN-inducible myxovirus resistant protein 1 (MxA), which enables the translocation of this cytokine into a trans-Golgi network (TGN)-mediated unconventional secretory pathway. Our study highlights a novel and synergistic pathway involving IFN and the NLRP3 inflammasome in SLE.

## Main

SLE is a highly heterogeneous disease. Thus, a diverse array of common and rare genetic variants have been reported, and clinical manifestations vary from mild skin and/or joint involvement to life-threatening disease^3^. Longitudinal studies assessing blood transcriptional pathways that track disease activity (DA) also support multiple pathogenic mechanisms, including activation of the Type I IFN, myeloid, B cell/plasma cell, lymphoid and/or erythroid pathways^4^. Although many SLE manifestations could be ascribed to excessive production of Type I IFN, blocking this pathway offers patial relief over standard of care in clinical trials, pointing to additional potential targets^5^.

Immune complexes (ICs) carrying NAs induce Type I IFN production in SLE^6,7^. In addition to NAs of nuclear origin, the role of mitochondrial NAs (mtNAs) has recently emerged. Indeed, internalization of SLE neutrophil-derived mtDNA by plasmacytoid dendritic cells (pDCs) triggers TLR9-dependent Type I IFN^8^. Furthermore, opsonization of Mito^+^ RBCs within macrophages (Mφ) leads to cGAS/STING-dependent production of Type I IFN^2^. In addition to TLR9 and cGAS, mtDNA is a ligand for pattern recognition receptors (PRRs) such as the NLRP3 inflammasome, which is involved in a variety of inflammatory diseases^9^. Activation of NLRP3, however, is not a feature of human interferonopathies perhaps due to the well-documented counter-regulation between the Type I IFN and IL-1β pathways^10^.

Here we identify erythrophagocytic Mo that co-express ISGs and IL-1β in the blood of SLE patients with active disease. This phenotype is recapitulated *in vitro* upon opsonization of Mito^+^ RBCs. Importantly, we identify a novel IL-1β secretory pathway dependent on the IFN-inducible protein MxA, which enables the entry of mature IL-1β (mIL-1β) into a TGN-mediated unconventional secretory pathway.

### Erythrophagocytic Mo co-expressing IL-1β and ISGs are expanded in the blood of patients with active SLE

Phagocytosis of Mito^+^ RBCs by Mφ stimulates the secretion of Type I IFN in a mtDNA- and cGAS-dependent manner^2^. Recent single-cell RNA sequence (scRNA-seq) studies of SLE PBMCs highlighted the presence of Mo co-transcribing IL1B and ISGs in patients with active disease^11^. As mtDNA is a ligand for both cGAS and the NLRP3 inflammasome^9^, we investigated the presence of erythroid markers, IL1β and ISGs in *ex-vivo* SLE blood Mo.

Classical (CD14^+^ CD16^-^) Mo were sorted and stained for MxA, a well-established ISG, and IL-1β. In healthy donors (HD), most cells (74.3% ± 10.8%) did not express either of these two proteins, while a fraction exclusively expressed either IL-1β (19% ± 9.9%) or MxA (6.6% ± 4.7%) (**Fig. S1a, S1b and S1c**). As expected, Mo isolated from patients with SLE or juvenile dermatomyositis (JDM), a disease that shares with SLE a prominent Type I IFN signature^12^, broadly expressed MxA (**Fig. 1a and S2a**). However, while JDM Mo did not co-express IL1β, up to ∼60% of SLE MxA^+^ Mo did (**Fig. 1a, S2a, S2b and Table S1**). Notably, IL-1β^+^ MxA^+^ Mo were not detected within intermediate (CD14^+^ CD16^+^) or non-classical (CD14^dim^ CD16^+^) Mo subsets (**Fig. S2c**).

**Fig. 1.**
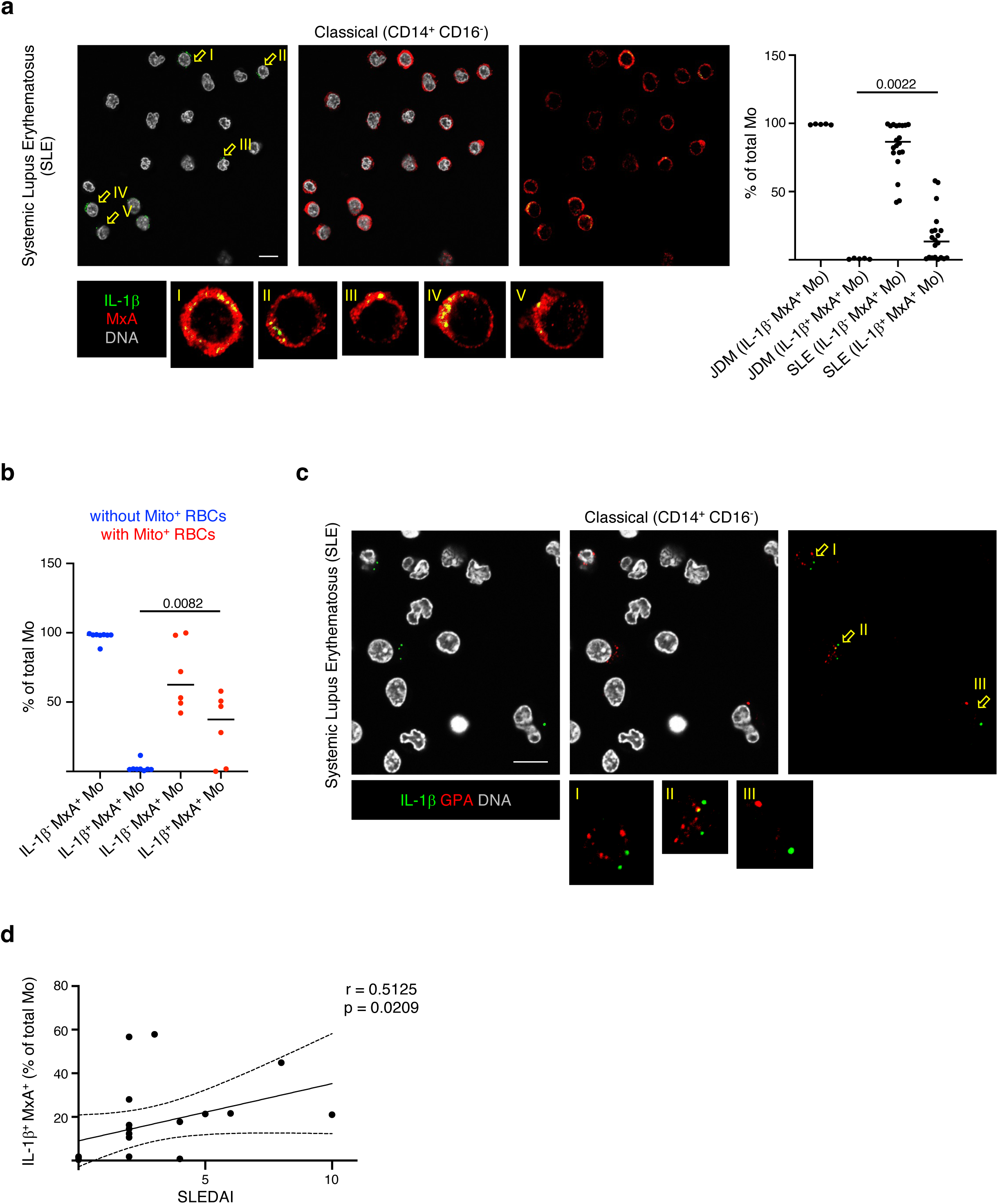
Erythrophagocytic Mo co-expressing IL-1β and ISGs are expanded in active SLE. **a,** Confocal images (left) and quantification (right) of classical Mo isolated from JDM and SLE patients. Each dot represents the average count from at least ten different micrscopy fields from a study participant sample that was processed, stained and analyzed independently. Scale bar: 20 µm. **b,** Percentage of IL-1β^−^ MxA^+^ Mo and IL-1β^+^ MxA^+^ Mo in SLE patients without (blue, n=8) or with (red, n=6) circulating Mito^+^ RBCs. **c,** Confocal images of classical Mo isolated from SLE patients showing evidence of erythrophagocytosis. GPA: Glycophorin A. Scale bar: 20 µm. **d,** Two-tailed Pearson’s correlation between SLEDAI and the percentage of IL-1β^+^ ISGs^+^ Mo (n=20). Dashed lines represent 95% confidence intervals. Each dot represents a study participant sample that was processed, stained and analyzed independently.

As IL-1β antibodies used in IF studies cannot discriminate mature IL-1β from its proform, we next explored whether IL-1β^+^ MxA^+^ SLE Mo released mIL-1β. Sorted Mo were placed on anti-mIL-1β-antibody-coated coverslips in order to capture mIL-1β released from individual cells while determining their viability with MitoTracker DeepRed (MTDR) staining^13^. Individual SLE Mo remained viable and released mIL-1β, demonstrating that *ex-vivo* SLE Mo are poised for mIL-1β production without undergoing cell death (**Fig. S3a**).

IL-1β^+^ MxA^+^ Mo were mainly detected in patients harboring circulating Mito^+^ RBCs (**Fig. 1b**), and ∼50% of them displayed cytosolic Glycophorin A (GPA) suggestive of erythrophagocytosis (**Fig. 1c and S3b**). Importantly, the levels of IL-1β^+^ MxA^+^ Mo, but not of IL-1β^-^ MxA^+^ Mo, directly correlated with DA according to the SLE disease activity index (SLEDAI)^14^ (**Fig. 1d, S3c, S3d and Table S1**).

Together, these data support that IL-1β^+^ MxA^+^ Mo include cells undergoing *in vivo* erythrophagocytosis and represent a better biomarker of SLE DA than ISG^+^ only Mo.

### RBC-derived mtNAs trigger Type I IFNs and IL-1β production in human Mo

To confirm that phagocytosis of Mito^+^ RBCs triggers the generation of IL-1β^+^ ISGs^+^ Mo, we used a cell culture model involving exposure of primed HD Mo to IgG-opsonized RBCs, which markedly increases erythrocyte internalization *in vitro* (**Fig. S4a**). Although Mito^-^ RBCs and Mito^+^ RBCs were internalized at similar rates, only the latter induced ISG upregulation (**Fig. 2a and S4b**). It is well known that Type I IFNs down-regulate inflammasome activity^10^. Strikingly, Mito^+^ RBC-treated Mo upregulated IL1B (**Fig. 2a**). Consistently, protein levels of both IP-10, a surrogate marker for activation of the IFN pathway^2,15^, and IL-1β were elevated in supernatants from Mito^+^ RBC-treated Mo (**Fig. 2b**). Immunoblotting MxA in the whole cell lysate (WCL) and IL-1β in the supernatant (Sup) confirmed these results **(Fig. S4c)**.

**Fig. 2.**
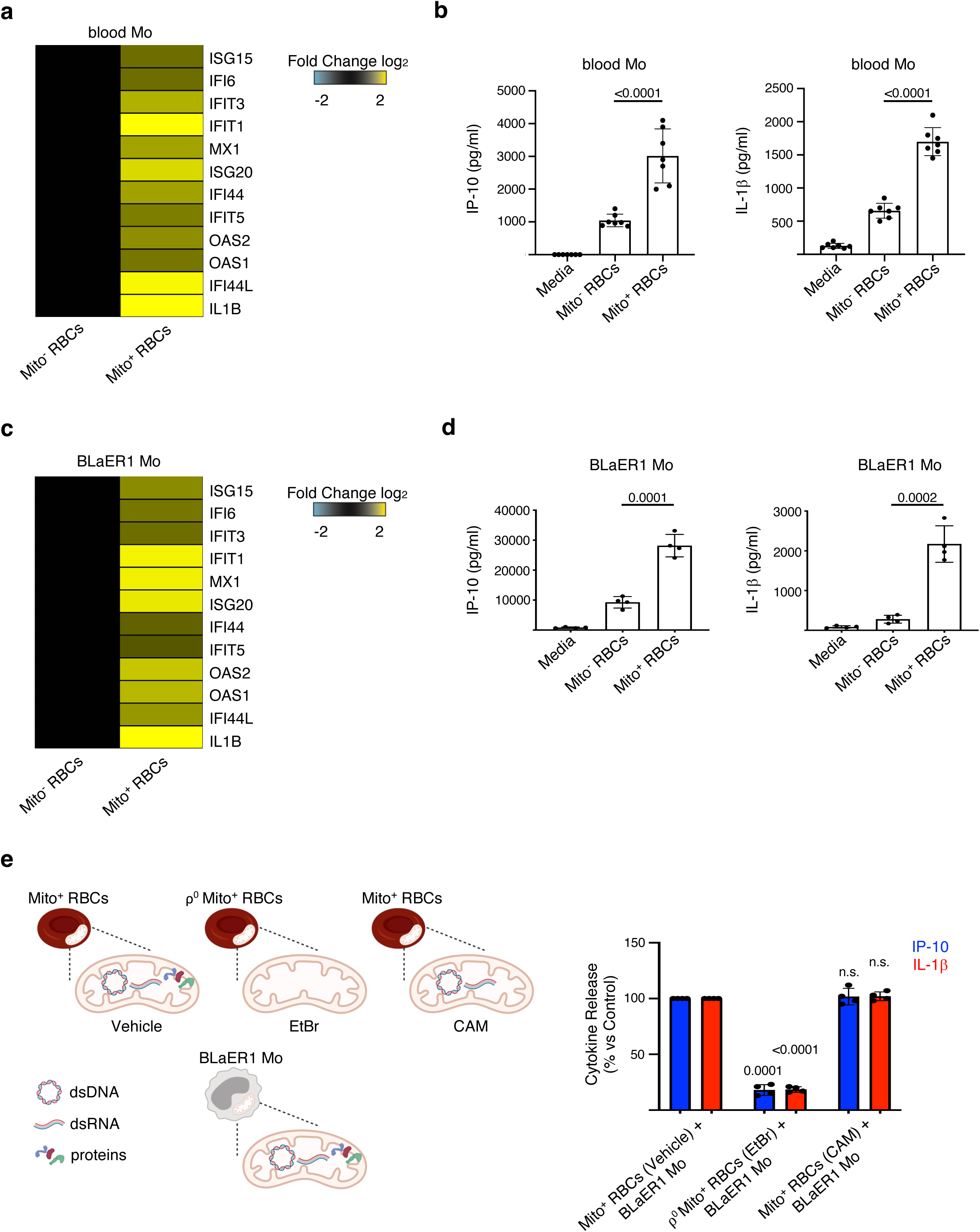
RBC-derived mtNAs trigger Type I IFNs and IL-1β production in human Mo. **a,** Heat map of differentially expressed genes in blood Mo that phagocytized either Mito^-^ or Mito^+^ RBCs. Data are normalized to Mito^-^ RBCs sample**. b,** Levels of IP-10 and IL-1β in the supernatants of blood Mo cultured with media, Mito^-^ or Mito^+^ RBCs. (n=6). **c,** Heat map of differentially expressed genes in BLaER1 Mo that phagocytized Mito^-^ or Mito^+^ RBCs. Data are normalized to Mito^-^ RBCs sample**. d,** Levels of IP-10 and IL-1β in the supernatants of BLaER1 Mo cultured with media, Mito^-^ or Mito^+^ RBCs. (n=4). **e,** Experiment scheme (left) and normalized cytokine levels (right) in the supernatants of BLaER1 Mo activated with Mito^+^ RBCs generated in the presence of vehicle, ethidium bromide (EtBr; ρ^0^) or chloramphenicol (CAM). (n=4).

Similarly, bone marrow (BM) Mo, which play a fundamental role in the RBC life cycle and participate in erythrophagocytosis^16^, responded to Mito^+^ RBCs by co-producing IP-10 and IL-1β (**Fig. S4d**). Thus, this phenotype is not SLE intrinsic but a response of myeloid cells to Mito^+^ RBCs.

To dissect the mechanisms underlying this dual cytokine production in a model system amenable to genetic manipulation, we turned to BLaER1 cells, a cell line that can be transdifferentiated into CD19^-^ CD11b^+^ BLaER1 Mo that fully recapitulate human Mo^15,17,18^ (**Fig. S4e**). Exposure of BLaER1 Mo to Mito^+^ RBCs recapitulated the results obtained with primary Mo, supporting that this cell line is a suitable model for our studies (**Fig. 2c, 2d, S4f and S4g**).

Mito^+^ RBCs are a source of mitochondrial damage-associated molecular patterns (mtDAMPs), which include mtNAs^2,19^. To assess whether Mito^+^ RBC-derived mtNAs were involved in Mo production of IP-10 and IL-1β, we generated Mito^+^ RBCs deficient in mtNAs (ρ^0^ Mito^+^ RBCs)^20^ (**Fig. S4h**). BLaER1 Mo exposed to ρ° Mito^+^ RBCs produced less IP-10 and IL-1β without major differences in erythrophagocytosis, suggesting that Mito^+^ RBC-derived mtNAs are the main drivers of this response (**Fig. 2e and S4i**). Since ρ° Mito^+^ RBCs lack not only mtNAs, but also mtDNA-encoded proteins (**Fig. S4j**), we treated Mito^+^ RBC cells with chloramphenicol (CAM) to selectively inhibit mitochondrial translation^20^ (**Fig. S4h and S4j**). Induction of both IP-10 and IL-1β was observed under these conditions (**Fig. 2e**), further implicating Mito^+^ RBC-derived mtNAs as major triggers of this type of Mo activation.

### Mito^+^ RBC-derived mtDNA directly activates cGAS but not NLRP3

Cytosolic mtDNA can activate the cGAS/STING pathway, which upon phosphorylation of TBK1 and IRF3 initiates a Type I IFN response^9^. Indeed, knockout (KO) of cGAS or STING reduced IP-10 secretion in response to Mito^+^ RBCs (**Fig. 3a and S5a**). Similar results were obtained upon pharmacological blockade of this pathway (**Fig. S5b**). Concurrent with the activation of STING, Mito^+^ RBC-treated Mo increased TBK1 and IRF3 phosphorylation (**Fig. S5c**).

**Fig. 3.**
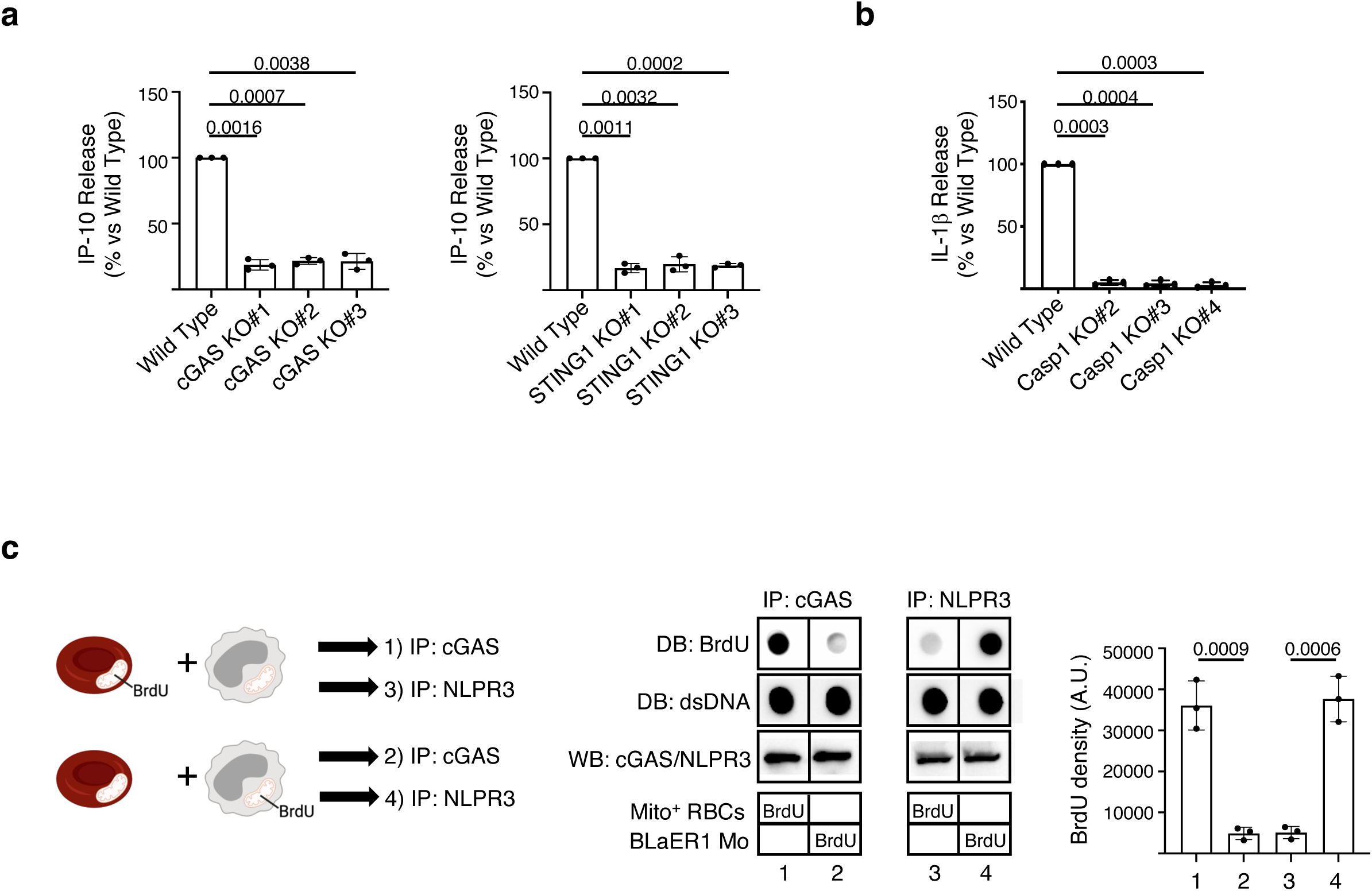
Mito^+^ RBC-derived mtDNA binds cGAS while Mo-derived mtDNA binds NLRP3. **a,** Normalized IP-10 levels in the supernatants of Mito^+^ RBCs-activated wild type, cGAS KOor STING1 KO BLaER1 Mo. (n=3). **b,** Normalized IL-1β levels in the supernatants of Mito^+^ RBCs-activated wild type or Caspase-1 KO BLaER1 Mo. (n=3). **c,** Experimental scheme (left), dot/Western blot analysis (middle) and quantification (left; n=3) of Mito^+^ RBCs (1 - 3) or BLaER1 Mo (2 − 4) labeled with bromodeoxyuridine (BrdU) and then immunoprecipitated (IP) with cGAS or NLRP3 antibodies. As a loading control, cGAS or NLRP3 immunoblot or double-stranded DNA (dsDNA) dot blot (DB) was performed. A.U.: Arbitrary Units.

Cytoplasmic dsDNA, including mtDNA, activates the AIM2 inflammasome in murine myeloid cells^21^. However, AIM2 is dispensable in sensing cytosolic dsDNA in human myeloid cells^15^. Accordingly, BLaER1 AIM2 KO Mo were still able to release IL-1β upon tranfection with dsDNA or internalization of Mito^+^ RBC (**Fig. S5d and S5e**). To test the extent to which IL-1β production was independent on AIM2 in human primary Mo, we adapted an antibody-targeting degradation approach (Trim-Away), to knockdown AIM2 protein^22^. The optimized Trim-Away protocol resulted in highly efficient delivery of antibody into the cytoplasm without cell death (**Fig. S5f and S5g**). Yet, supporting our results with BLaER1 Mo, the targeted reduction of AIM2 from human primary Mo did not affect the production of IL-1β induced by either tranfected dsDNA or Mito^+^ RBCs (**Fig. S5h**). Together, these data confirm that AIM2 is dispensable for Mito^+^ RBC-mediated inflammasome activation in human myeloid cells. Cytosolic mtDNA can also activate the NLPR3 inflammasome^23^. Indeed, IL-1β secretion was significantly reduced upon blockade of this pathway (**Fig. 3b, S5i and S5j**). Notably, reduced cytokine production was not a consequence of differential internalization of Mito^+^ RBCs, which was equivalent in wild type and KO BLaER1 Mo (**Fig. S5k**). Independent experiments with primary human Mo using specific inhibitors confirmed the key role of Mito^+^ RBC-derived mtNAs in the production of IP-10 and IL-1β through the activation of cGAS/STING and NLRP3/caspase-1, respectively (**Fig. S6a, S6b, S6c and S6d**).

As mtDNA binds to both cGAS and NLRP3^23,24^, we surmised that Mito^+^ RBC-derived mtDNA would be the ligand for these two sensors. Therefore, we immunoprecipitated (IP) cGAS or NLRP3 from BLaER1 Mo treated with Mito^+^ RBCs labeled with bromodeoxyuridine (BrdU), a thymidine analog that in post-mitotic cells selectively labels mtDNA^23^. BrdU signal was clearly detected in cGAS ICs, but labeled DNA did not co-immunoprecipitate (co-IP) with NLRP3, indicating that Mito^+^ RBC-derived mtDNA was not the NLRP3 ligand (**Fig. 3c**).

In human myeloid cells, a cGAS/STING-mediated lysosomal cell death (LCD) has been shown to be the default mode of NLRP3 activation in response to cytosolic dsDNA^15^. However, Mito^+^ RBCs did not elicit cytoplasmic translocation of cathepsin D, loss of plasma membrane (PM) integrity, or pyropotosome formation, all hallmarks of LCD-induced NLRP3 activation (**Fig. S7a, S7b, S7c and S7d**). Furthermore, pharmacological inhibition of LCD with Bafilomycin A1 (Baf A1) did not prevent Mito^+^ RBC-induced IL-1β secretion (**Fig. S7e**). LCD leads to a subsequent drop in cytoplasmic K^+^, which triggers NLRP3 activation^15^. Accordingly, blocking K^+^ efflux prevented IL-1β production upon stimulation with transfected dsDNA but not with Mito^+^ RBCs (**Fig. S7f**). Altogether, these data indicate that the cGAS-STING-LCD axis is not involved in NLRP3 activation in response to Mito^+^ RBCs.

### Mito^+^ RBC-derived mtRNA triggers the release of Mo-derived mtDNA fragments

As Mito^+^ RBC-derived mtDNA neither triggered IL-1β production through the STING-LCD pathway nor associated directly with NLPR3, we investigated alternative sources of NLRP3 stimulatory DNA. Thus, we surmised that Mito^+^ RBC phagocytosis might result in the cytosolic release of Mo-derived mtDNA. Indeed, NLRP3 IP from BrdU-labeled BLaER1 Mo exposed to unlabeled Mito^+^ RBCs enabled detection of BrdU signal (**Fig. 3c**). Accordingly, BLaER1 Mo lacking mtDNA^25^, but not mtDNA-encoded proteins, displayed defective IL-1β production upon internalization of Mito^+^ RBCs (**Fig. 4a, S8a and S8b**).

**Fig. 4.**
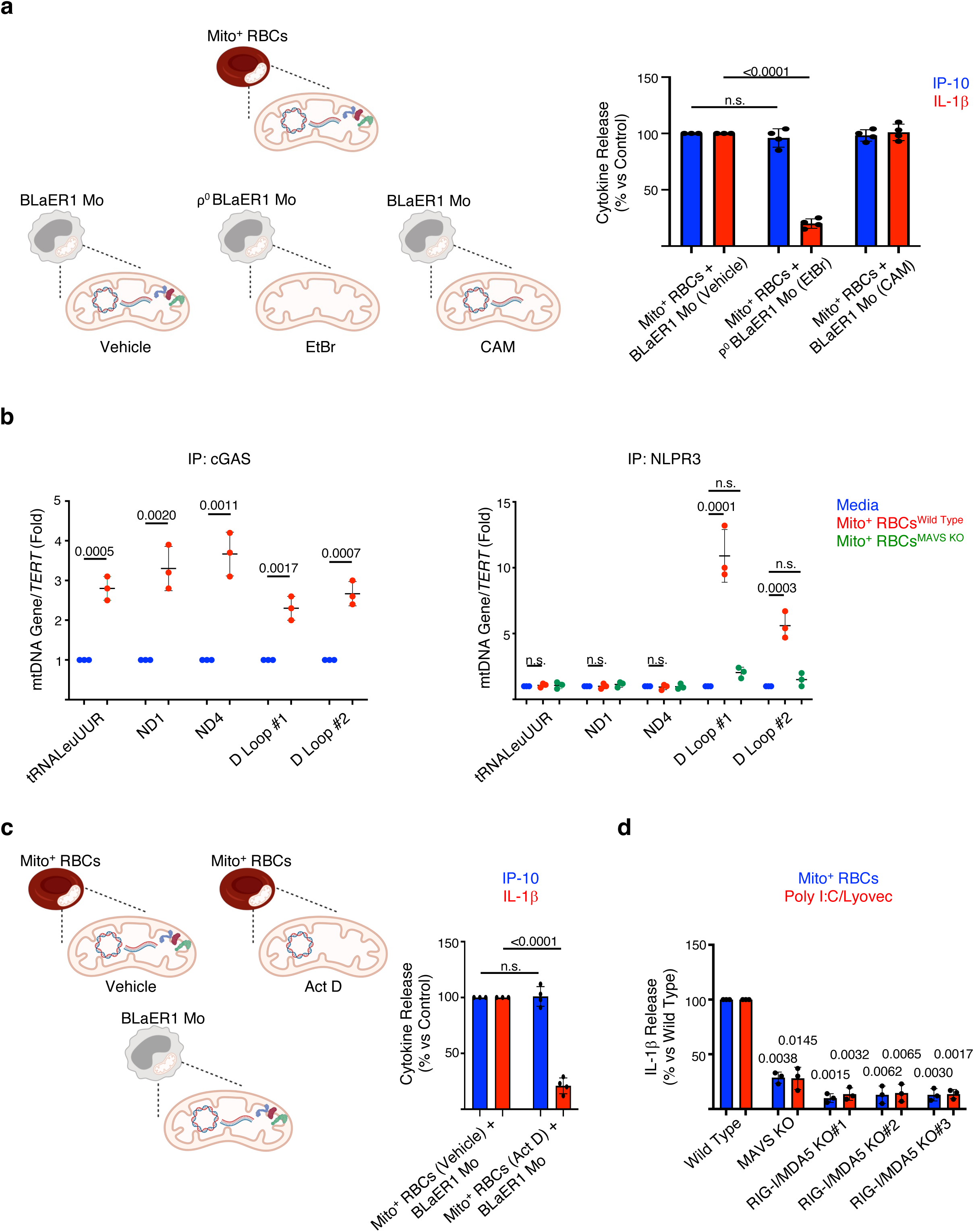
Mito^+^ RBC-derived mtRNA triggers the release of Mo-derived mtDNA fragments. **a,** Experiment scheme (left) and normalized cytokine levels (right) in the supernatants of BLaER1 Mo, generated in the presence of vehicle, EtBr (ρ^0^) or CAM and activated with Mito^+^ RBCs. (n=4). **b,** Relative levels of mtDNA-encoded genes in cGAS (left) or NLRP3 (right) immunocomplexes isolated from Mito^+^ RBC-activated wild type or MAVS KO BLaER1 Mo. (n=3). **c,** Experiment scheme (left) and normalized cytokine levels (right) in the supernatants of BLaER1 Mo cultured with Mito^+^ RBCs generated in the presence of Actinomycin D (Act D). (n=4). **d,** Normalized IL-1β levels in the supernatants of Mito^+^ RBCs- or Poly I:C/Lyovec-activated wild type, MAVS KO or RIG- I/MDA5 KO BLaER1 Mo. (n=3).

Our results show that Mito^+^ RBC-derived mtDNA preferentially binds cGAS while Mo-derived mtDNA binds to NLRP3, suggesting different immunostimulatory properties. Even though oxidation of deoxyguanosine to 8-hydroxy-2‘-deoxyguanosine (8-OHdG) increases the immunorecognition of DNA^8,23^, we did not findsignificant differences in 8-OHdG content of mtDNA associated with either cGAS or NLPR3 (**Fig. S8c and S8d**). Previous studies have shown that NLRP3 recognizes short dsDNA fragments, while cGAS requires larger fragments for efficient activation^18,23^. Thus, we extracted DNA from NLPR3 and cGAS ICs and subjected it to PCR amplification using primers flanking four mtDNA-encoded transcripts along the mitochondrial genome. While mtDNA associated with cGAS was enriched in all mtDNA transcripts tested, mtDNA bound to NLPR3 was only enriched in D-loop region transcripts (**Fig. 4b**). Accordingly, *in vitro* binding studies showed that cGAS and NLRP3 preferentially interacted with intact and fragmented mtDNA, respectively (**Fig. S8e**). Collectively, these results confirm the relevance of DNA length in the selective affinity of mtDNA for either cGAS or NLRP3.

We next explored the mechanism whereby Mito^+^ RBC-derived NAs indirectly trigger NLRP3 activation through the cytosolic release of Mo-derived mtDNA fragments. In addition to dsDNA, mitochondria carry immunogenic dsRNA^26,27^. Accordingly, Mito^+^ RBCs generated in the presence of Actinomycin D (Act D), which inhibits mtDNA transcription and thus reduces the levels of mt dsRNA^26^, failed to induce IL-1β production (**Fig. 4c**). In myeloid cells, cytoplasmic dsRNA activates NLRP3 in a MAVS-dependent manner, although the cytosolic sensor upstream of MAVS remains poorly understood^28,29^. Indeed, Mito^+^ RBC-derived mt dsRNA triggered the formation of MAVS oligomers, a hallmark of MAVS activation (**Fig. S8f**). Importantly, levels of mtDNA bound to NLPR3, activation of caspase-1, and secretion of IL-1β were greatly reduced in MAVS KO BLaER1 Mo (**Fig. 4b, 4d, S8g and S8h**). Similar results were obtained upon transfection of the synthetic dsRNA analog polyinosinic polycytidylic acid (poly I:C) (**Fig. 4d, S8i, S8j, S8k**). These results support a fundamental role of MAVS in the release of Mo-derived mtDNA fragments and NLRP3 activation induced by cytoplasmic dsRNA. dsRNA is sensed by cytoplasmic RLRs such as RIG-I and MDA5^28^. However, neither single RIG-I nor MDA5 KO significantly reduced IL-1β secretion in response to cytoplasmic dsRNA (**Fig. S9a and S9b**). Similarly, DHX33 KO^30^ did not affect IL-1β secretion (**Fig. S9c and S9d**). Instead, levels of mtDNA bound to NLPR3, activation of caspase-1, and secretion of IL-1β were greatly reduced in BLaER1 Mo deficient in both RIG-I and MDA5 (**Fig. 4d, S8h, S8k, S9e, S9f and S9g**), suggesting that these RLRs act redundantly in human Mo to sense cytoplasmic dsRNA, including mt dsRNA.

### MxA is required for unconventional secretion of mIL-1β

mIL-1β lacks the signal peptide for endoplasmic reticulum (ER)/Golgi-dependent secretion and is therefore released through non-canonical pathways, including passive release following pyroptosis^31^. Yet, caspase-1 mediated cleavage of gasdermin-D (GSDMD), whose N-terminal fragment (GSDMD-NT) subsequently forms pores in the PM, leads to pyroptosis and mIL-1β secretion^32^. However, neither Mito^+^ RBC nor Poly I:C induced GSDMD cleavage or lactate dehydrogenase (LDH) release, a hallmark of pyroptosis^33^ (**Fig. S10a and S10b**). GSDMD pores can also promote mIL-1β secretion from living cells independently of pyroptosis^32,33^. Yet, Mito^+^ RBC- or Poly I:C-activated BLaER1 Mo did not exhibit signs of pore formation, as assessed by propidium iodide (PI) influx^33^ (**Fig. S10b**). The amount of GSDMD-NT generated under these conditions could be below the antibody detection threshold and/or not be enough to cause detectable PI influx, yet it could be sufficient to enable membrane pore formation and mIL-1β release. However, mIL-1β secretion was not attenuated in GSDMD KO BLaER1 Mo, supporting that mIL-1β release in response to Mito^+^ RBCs or Poly I:C proceeds in a GSDMD-independent manner in human Mo (**Fig. S10c**).

Strikingly, IF analysis of SLE blood Mo revealed IL-1β co-localization with MxA but not with other ISGs such as ISG15 or IFIT1 (**Fig. 1a, S10d and S10e**). Co-localization between IL-1β and MxA was also observed in activated BLaER1 Mo (**Fig. S11a**). As MxA belongs to the dynamin-like large GTPase family, a group of proteins involved in endocytosis/exocytosis^34^, we surmised that it might be involved in the unconventional secretion of mIL-1β.

In humans, MxA and mIL-1β are cytosolic proteins, but they also interact with subcellular membranes^35,36^. Indeed, using differential centrifugation, we detected both proteins in the cytosol fraction (100k S) as well as in the microsomal pellet (100k P), while proIL-1β was restricted to the cytosol^37^ (**Fig. 5a and S11b**). We then resolved the microsomal pellet on a sucrose gradient and analyzed the distribution of MxA and mIL-1β in relation to different cellular compartments. Both MxA and mIL-1β were enriched in fractions 10 through 12, coincident with the TGN, and co-localized with this compartment by IF analysis (**Fig. 5b, S11c and S11d**). Within the 100k P fraction, mIL-1β but not MxA was resistant to proteinase K (PK) digestion (**Fig. S11e**), suggesting that MxA is on the surface while mIL-1β resides within the lumen of the TGN membranes.

**Fig. 5.**
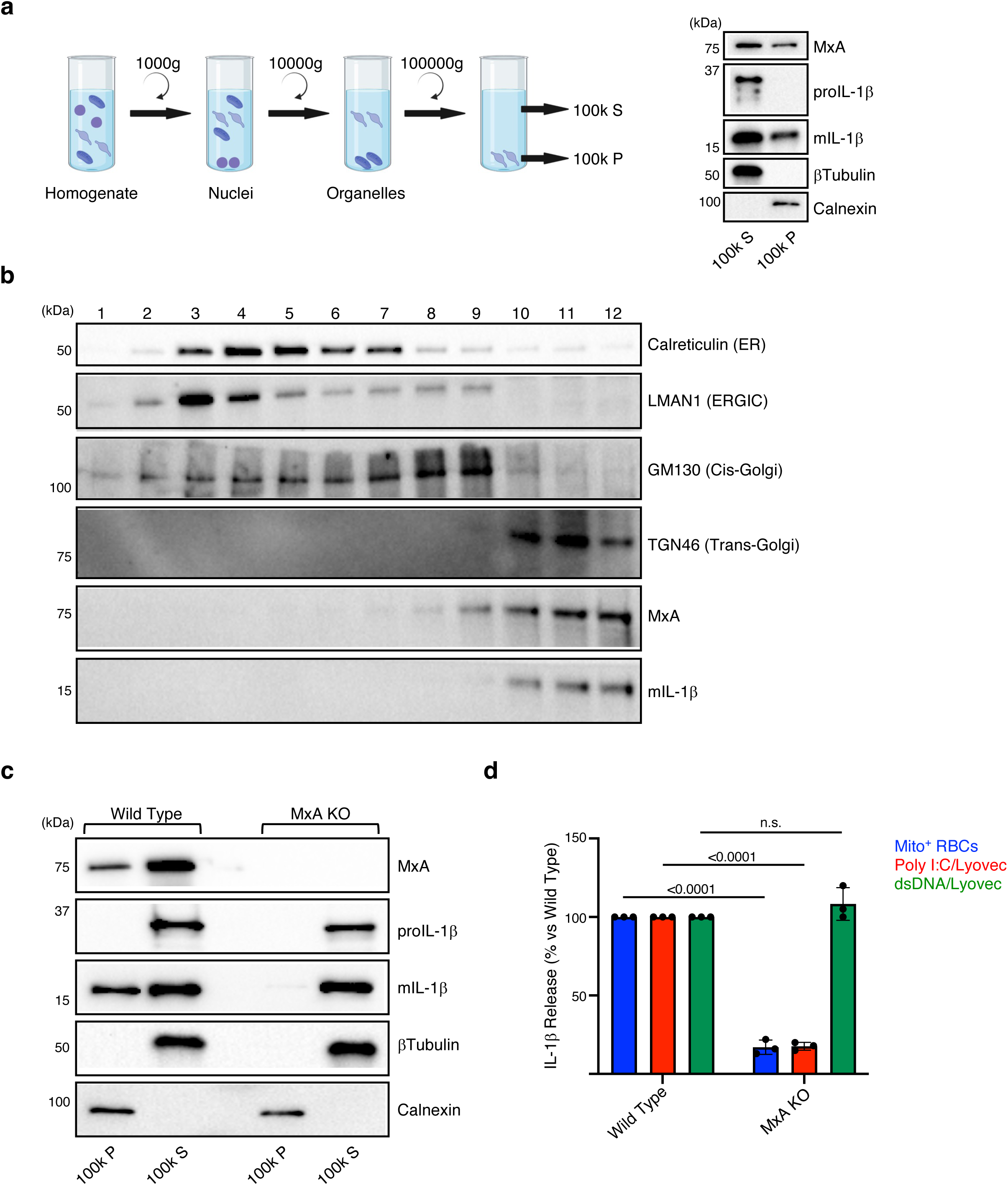
MxA is required for unconventional secretion of mIL-1β. **a,** Experimental scheme (left) and Western blot analysis (right) of the microsomal pellet (100k P) and postmicrosomal supernatant (100k S) isolated from Poly I:C/Lyovec-activated BLaER1 Mo. **b**, Western blot analysis of different fractions collected after sucrose fractionation of microsomal membranes isolated from Poly I:C/Lyovec-activated BLaER1 Mo. ER: endoplasmic reticulum; ERGIC: ER-Golgi intermediate compartment. **c,** Western blot analysis of the 100k P and 100k S fractions isolated from Poly I:C/Lyovec-activated wild type or MxA KOBLaER1 Mo. **d,** Normalized IL-1β levels in the supernatants from Mito^+^ RBCs, Poly I:C/Lyovec and dsDNA/Lyovec-activated wild type or MxA KO BLaER1 Mo. (n=3).

MxA forms dimers and tetramers in a criss-cross fashion and further oligomerizes into large structures^38^. Indeed, in activated cells, membrane-bound MxA formed SDS-resistant high molecular weight oligomers sensitive to reducing agents, suggesting that disulfide bond formation contributed to their stability (**Fig. S12a**). As mIL-1β interacts with and promotes the oligomerization of the ERGIC-resident protein TMED10^39^, we investigated whether this cytokine had similar effects on MxA. Indeed, MxA associated with endogenous mIL-1β, but not with pro-IL-1β (**Fig. S12b**). Furthermore, MxA oligomerization was blunted in BLaER1 Mo activated in the presence of a caspase-1 inhibitor, while it increased in IFN-primed cells transfected with recombinant mIL-1β (**Fig. S12a and S12c**). Notably, mIL-1β induced the formation of MxA oligomers without affecting the levels or the association of MxA with subcellular membranes (**Fig. S12d**). Finally, mIL-1β enhanced the oligomerization of recombinant MxA *in vitro* (**Fig. S12e**). Altogether, these data support that mIL-1β interacts and promotes the oligomerization of MxA.

MxA oligomerizes into ring-like structures that resemble protein channels^38,40^. We therefore surmised that MxA might mediate the translocation of mIL-1β across subcellular membranes. Indeed, MxA deficiency significantly decreased membrane-associated mIL-1β (**Fig. 5c and S12f**). Furthermore, the residual mIL-1β detected in MxA KO cell membranes was not protected from PK digestion, supporting the role of MxA in the translocation process (**Fig. S12f**). To further prove this, a proteoliposome (PtLp)-based transport assay was performed^39^ (**Fig. S12g**). MxA was incorporated within liposome (Lp) membranes by mixing purified MxA with detergent-solubilized Lp followed by detergent removal. A flotation assay, with and without urea, demonstrated the stable integration of MxA within the membranes, while a PK protection assay showed that >95% of MxA acquired a transmembrane localization (**Fig. S12h and S12i**). The presence of MxA in the PtLp membranes generated a fraction (∼50%) of mIL-1β protected from PK digestion, indicating that mIL-1β enters into the PtLp through MxA (**Fig. S12i**). These results were further confirmed by measuring the release of encapsulated mIL-1β from the lumen of Lp^33^, which showed that mIL-1β release was significantly enhanced when Lp were treated with recombinant MxA (**Fig. S13a**). Notably, the release of tetrameric LDH was minimal under these conditions indicating that MxA does not affect Lp integrity (**Fig. S13b**). These results collectively support that MxA promotes the translocation of mIL-1β across membranes.

As MxA plays a role in the incorporation of mIL-1β within the TGN/secretory pathway, we surmised that this ISG directly contributes to mIL-1β release. Accordingly, activation of MxA KO BLaER1 Mo with either Mito^+^ RBCs or transfected Poly I:C led to a significant reduction in mIL-1β secretion, which was overturned upon MxA re-expression (**Fig. 5d and S13c**). Notably, similar results were obtained in human primary Mo upon MxA knockdown using Trim-Away (**Fig. S13d**). To determine whether IL-1β maturation was sufficient to induce MxA-mediated secretion, we ectopically expressed mIL-1β in cells. As previously shown^37^, ectopically expressed mIL-1β was secreted at low levels in untreated cells (**Fig. S13e**). However, Type I IFN-mediated MxA upregulation boosted the secretion of mIL-1β g further indicating that MxA alone is sufficient to promote the release of this cytokine independently of inflammasome activation (**Fig. S13e**).

Overall, these data support a novel role for MxA in the unconventional secretion of mIL-1β by enabling the incorporation of this cytokine within a TGN/secretory pathway.

## Discussion

A variety of mechanisms, both genetic and/or stochastic, contribute to IFN production in SLE^41^. Among them, autoantibody-mediated internalization of Mito^+^ RBCs within Mφ triggers IFN production *in vitro*^2^. We have now identified SLE blood Mo carrying RBC remnants that co-produce IL-1β and ISGs. This phenotype is recapitulated upon opsonization of Mito^+^ RBCs within Mo *in vitro*. Importantly, we uncover a novel GSDMD- and pyroptosis-independent mIL-1β secretion pathway that requires MxA, a prototype ISG (**Fig. S14**).

Our data support that mtDNA from both the opsonized Mito^+^ RBCs and the phagocytic Mo contribute to this phenotype. Thus, Mito^+^ RBC-derived mtDNA triggers type I IFN production in Mo upon activating cGAS, while IL-1β production entails the association between NLRP3 and Mo-derived mtDNA fragments, which are released into the cytosol upon sensing of RBC-derived mtRNA (**Fig. S14**). Cytoplasmic dsRNA is known to induce MAVS oligomerization in order to activate the NLPR3 inflammasome, but the downstream pathways remained unresolved^28,29^. Our study shows that MAVS oligomerization leads to the cytosolic release of mtDNA fragments, which then bind and activate NLRP3. Upstream of MAVS oligomerization, the cytosolic sensor(s) responsible for recognition of mt dsRNA also remained controversial^26,27^. Our study reveals that, within human Mo, RIG-I and MDA5 act redundantly in this pathway. A similar finding had been previously shown in murine Mφ upon transfection with bacterial, but not mt, dsRNA^28^. These divergent results could be explained by different delivery routes that might affect mt dsRNA compartmentalization and interaction with RLRs. It is also plausible that phagolysosomal degradation of Mito^+^ RBCs induces the release of additional factor(s) synergizing with the sensing of mt dsRNA by RLRs. In this regard, metabolites released upon phagolysosomal digestion of *P. falciparum*-infected RBCs have been shown to amplify *P. falciparum* RNA sensing by endosomal TLRs^42^.

GSDMD-driven pyroptosis is considered the main mechanism involved in the release of mIL-1β. Increasingly, however, it is being noted that this cytokine can be secreted in the absence of cell death^33,37,43–45^. While mIL-1β release from live cells may still rely on the formation of GSDMD pores, growing evidence supports that GSDMD-independent mIL-1β release can also occur^17,37,43^. We identify a novel MxA/TGN-dependent pathway leading to mIL-1β secretion from live human Mo. We describe how mIL-1β triggers MxA oligomerization, which enables mIL-1β translocation into the TGN, a major secretory sorting station that directs newly synthesized proteins into different subcellular destinations, including the extracellular space^46^. This mechanism of MxA-mediated mIL-1β unconventional protein secretion (UPS) is reminiscent of another translocation process; the TMED10-channeled UPS (THU)^39^. In THU, mIL-1β interacts with TMED10 on the ERGIC and induces its oligomerization and formation of a protein channel. The TMED10 channel then translocates the cargo into the lumen of the ERGIC to be released^39^. It is enticing to speculate that, depending on the cell type and/or the nature of the inflammasome activator, these pathways promote GSDMD- and pyroptosis-independent secretion of mIL-1β . Notably, most laboratory mice strains carry nonfunctional MX1 alleles with either large deletions or nonsense mutations^47^, supporting the species-specificity of secretory pathways such as the one we here describe.

In addition to their well-recognized effects on immune activation, Type I IFNs have immunosuppressive functions through, among others, down-regulation of inflammasome activity^10^. Recent studies, however, suggest that in some pathological conditions, including atherosclerosis, metabolic diseases, and neuroinflammatory disorders, the inhibitory effect of Type I IFN on IL-1β production might be halted. Although the factors that prime cells for inflammasome signaling under these non infectious conditions are not known, endogenous triggers are most likely at play^48^. Indeed, a population of pro-inflammatory IL-1β^+^ ISG^+^ Ly6C^+^ Mo infiltrates the central nervous system (CNS) of aged mice^49^. Similarly, IL-1β^+^ IP-10^+^ Mo infiltrate the CNS of mice with experimental autoimmune encephalomyelitis (EAE), where they contribute to tissue damage^50^. In this scenario, IL-1β secretion might be important for Mo transmigration across tissues, such as the blood-brain barrier^51^. Furthermore, the production of IL-1β and IP-10 is a hallmark of human SLE blood monocytes capable of inducing *in vitro* alloreactions^52^. The fate of SLE IL-1β^+^ ISG^+^ Mo, including their potential to migrate to inflamed tissues where they might play a pathogenic role, will require further studies. Importantly, this novel pathway might represent a target for therapeutic intervention not only in SLE, but in other inflammatory scenarios where the counter-regulation between Type I IFN and the inflammasome is disrupted.

## Materials and Methods

### Human samples

Children and adolescents with SLE and JDM were enrolled in the Pediatric Rheumatology clinics at Texas Scottish Rite Hospital for Children and Children’s Medical Center in Dallas (TX). Study procedures were followed in accordance with protocols approved by the Institutional Review Boards at the University of Texas Southwestern Medical Center (IRB #22-0302461) and Weill Cornell Medical College (IRB #1711018757). Assent was obtained from patients between 7 and 17 years of age. Patients were evaluated by a standardized protocol during routine clinic visits every three months and more frequently if clinical symptoms warranted evaluation. Blood was collected in ACD tubes (BD Biosciences) and laboratory measurements were recorded. Clinical disease activity was assessed using the SLE Disease Activity Index (SLEDAI-2K). Relevant clinical information is depicted in **Extended Table 1**.

### Isolation of human primary monocytes (Mo)

For isolation of blood Mo, peripheral blood mononuclear cells (PBMCs) were isolated from healthy donor (HD) Leukopacks (New York Blood Center) with Lymphoprep gradient centrifugation. Mo were then isolated with EasySep Human Monocyte Enrichment Kit (Stem Cell Technologies) following the manufacturer protocol. Alternatively, classical (CD14^+^ CD16^-^), intermediate (CD14^+^ CD16^+^) or non-classical (CD14^dim^ CD16^+^) Mo subsets were FACS sorted from HD, SLE or JDM PBMCs with anti-CD16 (Clone #3G8) and anti-CD14 (Clone #M5E2) andibodies. For isolation of bone marrow Mo, CD14^+^ Mo were FACS sorted from frozen human bone marrow mononuclear cells (Stem Cell Technologies) on a BD FACS Melody Cell Sorted (BD Biosciences) equipped with BD FACSChorus software. Dead cells were removed with 7-Aminoactinomycin D (7-ADD, BD Biosciences).

### BLaER1 cells culture and differentiation

The BLaER1 Human B-cell Precursor Leukemia Cell Line was purchased from Millipore Sigma− Aldrich and cultured as described^53^. Briefly, BLaER1 cells were expanded in cRPMI medium (RPMI medium 1640 supplemented with l-glutamine, sodium pyruvate, HEPES buffer, penicillin-streptomycin and 10% v/v FBS). BLaER1 cells were trans-differentiated into Mo for 6-7 days in cRPMI medium containing 10 ng/mL of hrIL-3 (Stem Cell Technologies), 10 ng/mL hrM-CSF (Stem Cell Technologies) and 100 nM β-Estradiol (Sigma−Aldrich). The process of trans differentiation was checked using flow FACS analysis with anti-CD19 (Clone #HIB19) and anti-CD11b (Clone #ICRF44) antibodies on a Cytek Aurora Cytometer equipped with SpectroFlo Software (Cytek Biosciences). To generate ρ° BLaER1 cells, cells were cultured with 500 ng/ml of ethidium bromide (EtBr; Sigma−Aldrich) for 4 days and checked for mitochondrial and nuclear DNA content by RT-PCR. Where indicated, 10 µM of bromodeoxyuridine (BrdU, Thermo Fisher) was added to BLaER1 cells, at day 6 post-differentiation, for 24 h.

### Generation of stable knockout (KO) BLaER1 cells

A Cas9-stable BLaER1 cells line was generated by transducing BLaER1 cells with a lentiviral Cas9 nuclease expression vectors containing a human codon-optimized version of *S. pyogenes* Cas9 nuclease under the control of hCMV promoter and the fluorescent protein mKate2 (Horizon Discovery). 48 h after transduction, mKate2^+^ cells were FACS sorted, expanded, and validated for stable Cas9 expression by Western blot using anti-Cas9 antibody (Santa Cruz Biotech; sc-517386). Using the Neon Transfection System (Thermo Fisher), 2.5x10^5^ Cas9^+^ BLaER1 cells were transfected (1400 V − Width 20 − 1 pulse) with 4.7 mM of gene-specific gRNA (Horizon Discovery) and then cultured in 3 mL of cRPMI medium. 48 h after transfection, cells were sorted into 96-well plates (1 cell/well) using BD FACS Melody Cell Sorted (BD Biosciences). Two to three weeks after sorting, 12 to 14 clones were selected and expanded for 96 h before assessing knockout efficiency by Western blot.

### Plasmid generation

DNA inserts containing the coding DNA sequence (CDS) for each wild type gene were generated by restriction digestion previous amplification from plasmids obtained from Addgene (h*MX1*: plasmid #158640; h*IL1B*: plasmid #166783) using primers flanked by NheI and BstBI restriction sites. Inserts were introduced into the pCDH-CuO-MCS-EF1a-CymR-T2A-Puro SparQ™ All-in-one Cloning and Expression Lentivector (QM800A-1, System Biosciences). The correct sequence of the gene of interest (GOI) in all generated plasmids was checked by Sanger sequencing. List of primers used *MX1* WT FWD (AAAAGCTAGCACCatggttgtttccgaagtggaca); *MX1* WT RV (TTTTTTCGAACTAaccggggaactgggcaa); *IL1B* WT (117-269) FWD (AAAAAAGCTAGCcaccatggcacctgtacgatcactgaactgc); *IL1B* WT (117-269) RV (TTTTTTTTCGAActaggaagacacaaattgcatggtgaagtcagttatat).

### Lentiviral transduction of BLaER1 cells

VSV-G pseudotyped lentiviral vector particles containing the GOI were generated in HEK 293T cells using a third-generation lentiviral packaging system for further infection of BLaER1 cells and delivery of the GOI. Work with lentiviral particles was exclusively carried out under S2 biosafety conditions. 0.5 x10^6^ HEK 293T cells were seeded in a well of a 6-well plate and incubated overnight in a humidified incubator at 37 °C with 5% CO2. The medium of the HEK 293T cells was exchanged with 1 mL fresh complete DMEM and cells were transiently co-transfected with four plasmids: 0.25 µg of a plasmid encoding the retroviral structure Gag-Pol proteins (pMDLg/pRRE, Addgene plasmid #12251), 0.5 µg of a plasmid encoding the vesicular stomatitis virus G protein (pMD2.G, Addgene plasmid #12259), 0.25 µg of a plasmid encoding for the transactivating protein Rev (pRSV-Rev, plasmid #12253) and 1.5 µg of a lentiviral vector plasmid that encodes the GOI. 24 h after transfection, the medium of the HEK 293T cells was exchanged to 2 mL fresh complete RPMI containing 30% FCS. 24 h later, the virus-containing supernatant of the HEK 293T cells was collected and filtered through a 0.45 µm filter to exclude cellular debris. This virus-containing supernatant was added to 0.1 x106 BLaER1 cells with 2 µg/mL polybrene and were spin-transduced by centrifugation at 800 x g for 45-60 min at 32°C. After centrifugation, cells were resuspended and seeded in a 24-well plate. After 24 h incubation with the viral particles, the virus-containing supernatant was removed and exchanged to fresh complete RPMI. Cells were incubated under S2 biosafety conditions for three more passages before they could be used under S1 biosafety conditions in further experiments. After the third passage, positively transduced cells were selected with 4 µg/mL puromycin for 7 days.

### Mito^+^ RBC generation

Mito^+^ RBCs were generated as previously described^2^. Briefly, PBMCs were seeded at 1×10^7^ cells/mL in expansion medium (phase 1), consisting of StemSpan Serum-Free Expansion Medium (SFEM, Stem Cell Technologies) supplemented with SCF (100 ng/mL; Stem Cell Technologies), PROCRIT (2 U/mL; Janssen), dexamethasone (1 µM; Sigma−Aldrich), IGF-1 (40 ng/mL; Stem Cell Technologies), cholesterol-rich lipids (40 µg/mL; Sigma−Aldrich) and IL-3 (10 ng/mL; Stem Cell Technologies). After 5 days, erythroblasts were purified by density purification using Lymphoprep and a lineage depletion step (Lineage Cell Depletion Kit; Miltenyi Biotech) was performed to ensure complete removal of lineage positive cells at this stage. Cells were reseeded in expansion medium without IL-3 (expansion phase - phase 2) and kept between 1.5 and 2×10^6^/mL by daily counting and medium changes. After 8 days of expansion cells were washed with phosphate-buffered saline (PBS) and reseeded in differentiation medium consisting of IMDM medium (differentiation phase - phase 3) supplemented with PROCRIT (10U/mL; Janssen), insulin (10 µg/mL; Santa Cruz Biotech), Human Peripheral Blood Plasma (3%; Stem Cell Technologies), 3,3‘,5-Triiodo-L-thyronine (1 µM; Sigma−Aldrich−Aldrich), IGF-1 (40ng/mL; Stem Cell Technologies) and holo-transferrin (0.5 mg/mL; Santa Cruz Biotech). Cells were kept between 2−4×10^6^ cells/mL by daily counting and medium changes. To generate Mito^+^ RBCs, MG132 (5 mM; Santa Cruz Biotech) was added to the culture during phase 3. To generate ρ° Mito^+^ RBCs, cells were cultured with MG132 (5 mM; Santa Cruz Biotech) and 500 ng/mL of ethidium bromide (EtBr; Sigma−Aldrich) during phase 3 and checked for mitochondrial and nuclear DNA content by RT-PCR. Alternatively, 100 µM of chloramphenicol (CAM; Santa Cruz Biotech), 0.5 mg/mL of Actinomycin D (Act D; Santa Cruz Biotech) or 10 µM of BrdU (Thermo Fisher) were added to the culture, at day 6 post-differentiation, for 24 h.

### Cell stimulation

Cells were primed with 2 µg/mL (human primary Mo) or 10 µg/mL (BLaER1 Mo) of Pam3CSK4 (Invivogen) for 2 h before activation. Mito^+^ and Mito^-^ RBCs, opsonized with anti-human RBC antibody (1:250; Rockland), were then added to primed Mo at a 1:2 (Mo to RBCs ratio) in cRPMI medium. Alternatively, primed Mo were stimulated with 200 ng of dsDNA from herring testes (dsDNA; Sigma−Aldrich) complexed with Lyovec (Invivogen), 500 ng of Poly I:C (Invivogen) complexed with Lyovec (Invivogen), or 2 μM of Nigericin (Sigma). Alternatively, Mo were primed with 200 U/mL human recombinant IFNα2β (Intron A; Merck) for 2 h and then stimulated with 20 ng/mL of recombinant human mIL-1β (R&D) complexed with Lyovec (Invivogen). Activation was carried out for 18 h unless stated differently. For BLaER1 Mo expressing inducible wild type MxA or mIL-1β, differentiated BLaER1 Mo were stimulated, 4 h before activation, with 1x Cumate solution (System Biosciences) to induce protein expression. Cells were then treated with Mito^+^ RBCs (MxA expressing cells) or with 200 U/mL human recombinant IFNα2β (mIL-1β expressing cells). For small-molecule inhibition, compounds were added 2 hr before stimulation of cells at the following concentrations: 5 µM MCC950 (Invivogen), 10 µg/mL Ac-YVAD-cmk (Invivogen), 5 µg/mL RU.521 (Invivogen), 5µM H-151 (Invivogen) or 50 nM Bafilomycin A1 (Invivogen). For blocking the efflux of K^+^, cells were incubated in medium that was diluted with 135 mM KCl (Roth) in sterile water containing 10% FCS to a final [K^+^] of 50 mM. The supernatants were collected and frozen for the quantification of cytokine levels with the Flex Set Cytometric Bead Array (CBA; BD Bioscience) using the supplier’s protocol.

### Quantification of mtDNA abundance

The abundance of mtDNA was assessed by Real-Time PCR as described previously^8^. Briefly, total DNA was extracted with DNAzol Reagent (Thermo-Fisher). DNA concentration was assessed with Quant-iT PicoGreen dsDNA Assay Kit (Thermo-Fisher). Real-Time PCR was performed with Power SYBR Green PCR Master Mix (Thermo-Fisher) with 3 ng of isolated DNA and 0.5 µM of the following primers: mtDNA encoded NADH dehydrogenase subunit 1 (ND1; 5‘-GCATTCCTAATGCTTACCGAAC-3‘ and 5‘-AAGGGTGGAGAGGTTAAAGGAG-3‘); genomic DNA encoded Glyceraldehyde 3-phosphate dehydrogenase (GAPDH; 5‘-AGGCAACTAGGATGGTGTGG-3‘ and 5‘-TTGATTTTGGAGGGATCTCG-3‘).

### Immunofluorescence (IF) analysis

For human primary Mo, sorted cells (0.5x10^5^ cells) were settled on poly-L-lysine coated glass coverslips for 10 min and then fixed with 4% paraformaldehyde for 20 min at room temperature. For BLaER1 cells, 2.5x10^5^ cells were differentiated into Mo on poly-L-lysine coated glass coverslips for 6 days. After 6 days, the differentiation medium was replaced with fresh medium containing 10 µg/mL of Pam3CSK4 (Invivogen). After 2 h, 500 ng of Poly I:C (Invivogen) complexed with Lyovec (Invivogen) was added. After 6 h cells were fixed with 4% paraformaldehyde for 20 min at room temperature. Cells were permeabilized with 0.01% Triton X-100 in PBS for 5 min at room temperature and then treated with BlockAid Blocking Solution (Thermo Fisher) supplemented with FcR Blocking Reagent (Miltenyi) for 30 min at room temperature. Staining was performed by using anti-IL-1β (1 µg/mL; R&D #MAB201-100), anti-Glycophorin A (GPA; 1 µg/mL; Biolegend #349102), anti-IFIT1 (1 µg/mL; Thermo Fisher #PA3-848), anti MxA (1 µg/mL; Abcam #ab95926), anti-ISG15 (1 µg/mL; Thermo Fisher #15981-1-AP) and anti-TGN46 (1 µg/mL; SantaCruz #sc-166594 or Thermo Fisher #PA5-23068) antibodies diluted in BlockAid Blocking Solution (Thermo Fisher). Isotype specific anti−mouse, anti−rabbit or anti-goat Alexa Fluor 488, 568 or 633 (Thermo-Fisher) were used as secondary antibodies. Nuclei were stained with Hoechst 33342 (Thermo Fisher) and samples were mounted with ProLong Gold Antifade Mountant (Thermo Fisher). The samples were acquired with a Zeiss LSM 880 confocal laser-scanning microscope equipped with a 63x/1.4 oil objective and ZEISS ZEN microscope software. Fiji/ImageJ software was used for analysis.

### Visualization of mIL-1β secretion from single cells

This method was adapted from Cullen et al.^13^. Briefly, poly-L-lysine coated coverslips were coated overnight with 4 µg/mL of human mIL-1β capture antibody (from R&D Human IL-1 beta/IL-1F2 DuoSet ELISA) in PBS. Sorted Mo (CD14^+^ CD16^-^) were then seeded overnight on mIL-1β capture antibody coated-coverslips in cRPMI medium supplemented with 10 ng/mL hrM-CSF (Stem Cell Technologies). 30 min before harvesting Mitotracker DeepRed (MTDR; Cell Signaling) was added at a concentration of 250 nM. Human mIL-1β detection antibody (from R&D Human IL-1 beta/IL-1F2 DuoSet ELISA) was added at a 0.2 µg/mL to the cells on the coverslips for 2 h at room temperature. Streptavidin conjugated Alexa Fluor 568 (Thermo Fisher) was added at a dilution of 1:2000 in PBS for 1 h at room temperature. Following gently washing with PBS, the coverslips were fixed in 4% paraformaldehyde. Nuclei were stained with Hoechst 33342 (Thermo Fisher) and samples were mounted with ProLong Gold Antifade Mountant (Thermo Fisher). The samples were acquired with a Zeiss LSM 880 confocal laser-scanning microscope equipped with a 63x/1.4 oil objective and ZEISS ZEN microscope software. Fiji/ImageJ software was used for analysis.

### SDS-PAGE and Western blot

Cultured cells were washed in PBS, and then lysed with RIPA buffer in the presence of Halt Protease and Phosphates Inhibitor Cocktail (Thermo Fisher). Samples were incubated on ice for 30 min and then centrifuged (13,000 g for 10 min at 4°C). The lysate containing the protein fraction were collected and stored at −80°C until further analysis. Protein concentration was estimated using the BCA kit (Thermo Fisher) following the manufacturer’s instructions. 20-30 µg of proteins were resuspended in 5× Lane Marker Reducing Sample Buffer (Thermo Fisher), boiled for 5 min at 100°C, and then subjected to electrophoresis with Mini-PROTEAN TGX Precast Gel (Bio-Rad Laboratories). Proteins were then transferred to nitrocellulose membranes, blocked for 1 h in 5% nonfat dry milk in TBST (Tris Buffered Saline containing 0.1% Tween-20), and incubated overnight at 4°C with the following primary antibodies: anti-MxA (Abcam #ab95926), anti-MTCO2 (Abcam #ab110258), anti-βactin (Abcam #ab8226), anti-phospho IRF3 (Cell Signaling #4947S), anti-phospho TBK1 (Cell Signaling #5483S), anti-NLRP3 (AdipoGen #AG-20B-0014-C100), anti-Caspase 1 (Cell Signaling #24232S), anti-polγ (Abcam #ab128899), anti-GAPDH (Abcam #ab181602), anti-COXIV (Abcam #ab202554), anti-cGAS (Cell Signaling #15102S), anti-STING (Cell Signaling #13647S), anti-cathepsin D (Abcam #ab75852), anti-MAVS (Cell Signaling #3993S), anti-RIG-I (Cell Signaling #3743S), anti-MDA5 (Cell Signaling #5321S), anti-DHX33 (Abcam #ab182006), anti-GSDMD NT (Cell Signaling #36425S), anti-cleaved (mature) IL-1β (Cell Signaling #83186S), anti-Calnexin (Cell Signaling #2679S), anti-Calreticulin (Cell Signaling #12238S), anti-LMAN1 (Novus Bio #NBP2-03381), anti-GM130 (Novus Bio #NBP2-53420SS), anti-TGN46 (SantaCruz #sc-166594), anti-IL-1β (R&D #MAB201-100), anti-cleaved (mature) Caspase 1 (Cell Signaling #4199S), anti-AIM2 (Cell Signaling #12948S), anti-ASC (SantaCruz #sc-514414), anti-GSDMD (Cell Signaling #97558S), anti-HSP90 (Cell Signaling #4877T) and anti-V5 Tag (Cell Signaling #13202S). After washing in TBST, the membranes were incubated for 1 h at room temperature with Poly HRP-conjugated anti−rabbit or anti−mouse IgG (Thermo Fisher). ECL Plus reagents (GE Healthcare) were used for detection. Digital images were acquired with ChemiDoc MP System (Bio-Rad Laboratories) and analyzed with Image Lab Software (Bio-Rad Laboratories). For measurement of mIL-1β in the supernatant by Western blot, proteins in the supernatant were precipitated with 1% (v/v) StrataClean resin (Agilent Technologies) overnight at 4°C. The resin was then treated with 5× Lane Marker Reducing Sample Buffer (Thermo Fisher). To generate the Triton X-100-soluble and -insoluble fractions, cells were lysed with 50 mM Tris-HCl (pH 7.6) containing 0.5% Triton X-100, EDTA-free protease inhibitor cocktail and phosphatase inhibitor cocktail (Thermo Fisher). The lysates were centrifuged at 6,000 g at 4°C for 15 min, and the pellets and supernatants were used as the Triton-insoluble and -soluble fractions, respectively.

### Immunoprecipitation (IP) and co-immunoprecipitation (co-IP)

Cells were washed twice with PBS and IP was performed using the Pierce Classic Magnetic IP/Co-IP Kit (Thermo Fisher) per manufacturer’s instructions. In brief, cells were activated as described above and lysed in IP lysis buffer (Thermo Fisher) supplemented with Protease Inhibitor Cocktail (Thermo Fisher), incubated with anti-cGAS (sc-515777; Santa Cruz Biotech), anti-NLPR3 (AdipoGen #AG-20B-0014-C100) or with V5-Trap® Magnetic Agarose (Proteintech). Samples were then rotated overnight at 4°C. For cGAS, NLRP3 and IL-1β IP, agarose magnetic beads were added and incubated with lysates on a rotator for 1 h at room temperature. Beads were then washed, and bound fractions were eluted with 5× Lane Marker Reducing Sample Buffer (Thermo Fisher). For detection of BrdU, dsDNA and 8-OHdG in the IP products, the eluted samples were dot-blotted and UV cross-linked to a nitrocellulose membrane that was immunoblotted with anti-BrdU (BioRad #AHP2405), anti-8OHdG (Thermo Fisher #BS-1278R) or anti-dsDNA (Novus Bio #NBP3-07302) antibodies. For the detection of the proteins in the IP products the eluted samples were heated to 95°C for 5 min, the gel was separated by SDS−PAGE, transferred to a PVDF membrane and immunoblotted with antibodies against NLRP3 (Cell Signaling #15101S), cGAS (Cell Signaling #15102S), V5 Tag (Cell Signaling #13202S), IL-1β (R&D #MAB201-100), mIL-1β (Cell Signaling #83186S), GAPDH (Abcam #ab181602), HSP90 (Cell Signaling #4877T) or MxA (Abcam #ab95926). For the detection of mtDNA in the immunoprecipitation products, DNA was extracted from eluted samples and qPCR was performed to amplify different regions of the mitochondrial genome. Nuclear DNA encoding TERT was used for normalization. Primer sequences are as follows: D loop #1 (F: CCAGTCTTGTAAACCGGAGA; R: CTATCACCCTATTAACCACTC), D loop #2 (F: CAGTCAAATCCCTTCTCGTC; R: TCCTTTTGATCGTGGTGATT), tRNALeuUUR (F: AGGACAAGAGAAATAAGGCC; R: CACGTTGGGGCCTTTGCGTA), ND1 (F: GGCTATATACAACTACGCAAAGGC; R: GGTAGATGTGGCGGGTTTTAGG), ND4 (F: CCCTCGTAGTAACAGCCATTCTC; R: CGACTGTGAGTGCGTTCGTAGT), TERT (F: GCCGATTGTGAACATGGACTACG, R: GCTCGTAGTTGAGCACGCTGAA). To assess the affinity of intact (Int^mtDNA^) or fragmented (Fgt^mtDNA^) mtDNA for cGAS and NLRP3, mtDNA was extracted from 2×10^7^ of Mito^+^ RBCs with the Mitochondrial DNA Isolation Kit (Abcam) and quantified with Quant-iT PicoGreen dsDNA Assay Kit (Thermo Fisher). mtDNA was biotinylated with Label IT Nucleic Acid Labeling Kits (Mirus) and fragmented with HaeIII (New England Biolabs) per manufacturer’s instructions. The efficiency of biotinylation and fragmentation was evaluated by Dot Blot and agarose gel electrophoresis respectively. BLaER1 cells were primed with 10 µg/mL of Pam3CSK4 (Invivogen) for 2 h and lysed in IP lysis buffer (Thermo Fisher) supplemented with Protease Inhibitor Cocktail (Thermo Fisher). 100 mg of IP lysate was incubated with 6 mg of biotinylated WT^mtDNA^ or Int^mtDNA^ for 2 h at 4°C followed by incubation, with anti-cGAS or anti-NLPR3 antibodies, overnight at 4°C. Magnetic beads were added and incubated with lysates on a rotator for 1 h at room temperature. Beads were then washed, and bound fractions were eluted with 5× Lane Marker Reducing Sample Buffer (Thermo Fisher). For detection of biotynilated DNA in the IP products, the eluted samples were dot-blotted and UV cross-linked to a nitrocellulose membrane that was immunoblotted with Pierce High Sensitivity Streptavidin-HRP (1:2500; Thermo Fisher).

### In vitro MxA oligomerization assay

1 µg/mL of purified human MxA (Novus Biological) was incubated with or without 100 ng/mL of recombinant human mIL-1b (R&D) for 30 min at room temperature in 20 mM HEPES buffer pH 7.4 and then incubated for 15 min with the cross-linking reagent BS(PEG)5 (1 mM). Zeba Spin Desalting Columns (7 K MWCO, 2ml. Thermo Fisher) were then used to separate crosslinked proteins from excess crosslinker and reaction byproducts. Samples were then subjected to SDS-PAGE and immunoblotting as described above.

### Cell fractionation and sucrose gradients

8×10^7^ BLaER1 cells were lysed in 1 mL of 25 mM HEPES pH7.5, 2.5 mM MgCl2, 50 mM Sodium acetate, 1 mM DTT and Protease Inhibitor Cocktail (Thermo Fisher). The lysates were disrupted using a Dounce homogenizer and centrifuged at 1,000 g for 5 min at 4°C. The resulting postnuclear supernatant was centrifuged first at 10,000 g (10 min at 4°C) and then 100,000 g (30 min at 4°C) to obtain 100k S and 100k P. Pellets were resuspended in 5× Lane Marker Reducing Sample Buffer (Thermo Fisher) and analyzed by Western blot. For flotation analysis, the 100k P membrane pellet was suspended in 1 mL 2.1 M sucrose buffer and layered below 10 mL of a 1.9-1.1 M discontinuous sucrose gradient. The gradient was subjected to centrifugation at 150,000 g for 18 h in a Beckman SW 40 rotor at 4°C and fractions of 500 ml were collected from the top of the gradient as described^54^. Proteins were extracted from sucrose gradient fractions by incubation with 1% (v/v) StrataClean resin (Agilent Technologies) overnight at 4°C, followed by centrifugation at 850 × g for 5 min at 4°C, aspiration of supernatant and resuspension in 5× Lane Marker Reducing Sample Buffer (Thermo Fisher). Samples were then subjected to SDS-PAGE followed by immunoblot analysis with the indicated antibodies. The cytosol fraction was isolated from 2×10^7^ BLaER1 cells using the Mitochondria Isolation kit (Thermo Fisher) following the manufacturer’s instructions. For SDS-PAGE, proteins were extracted from the cytosol fraction by incubation with 1% (v/v) StrataClean resin (Agilent Technologies) overnight at 4°C, followed by centrifugation at 850 × g for 5 min at 4°C, aspiration of supernatant and resuspension of the pellet in 5× Lane Marker Reducing Sample Buffer (Thermo Fisher). Samples were then subjected to SDS-PAGE followed by immunoblot analysis with the indicated antibodies.

### Semidenaturing detergent agarose gel electrophoresis (SDD-AGE)

SDD-AGE was performed according to a published protocol with minor modifications^55^. Briefly, the mitochondrial pellet (for MAVS oligomerization) or the 100k P membrane pellet (for MxA oligomerization) was resuspended in sample buffer (0.5× tris-borate EDTA, 10% glycerol, 2% SDS, and 0.0025% bromophenol blue) with or without β-mercaptoethanol (bME; 35 mM) and loaded onto a 1.5% agarose gel prepared with 1× T tris-borate EDTA, 0.1% SDS. After electrophoresis in running buffer (1× T tris-borate EDTA, 0.1% SDS) for 1 h with a constant voltage of 75 V at 4°C, proteins were transferred to PVDF membranes with a Trans-Blot Turbo Transfer System (Biorad) in preparation for Western blotting analysis.

### Flow cytometry analysis

Mitochondrial content in mature RBCs was assessed as previously described^2^. Briefly, PBMCs were removed from peripheral blood by density centrifugation using Lymphoprep™ (Stem Cell Technologies). The cell pellet was washed four times (3000 rpm, 8 min, 4°C) with PBS supplemented with 2 mM EDTA and 10 mM Glucose. The cell pellet was then re-suspended in PBS supplemented with 10 mM Glucose and MitoTracker Deep Red (MTDR, 200 nM; Thermo-Fisher) and incubated for 30 min at 37°C. The cell pellet was then washed four times with PBS, treated with Fc Block (BD Bioscience) and stained with anti-CD235a and anti-CD71 antibodies. The fluorescence intensity of MTDR was measured, in the mature CD71^-^ CD235a^+^ RBCs population, by flow cytometry on a BD FACS Canto II (BD Biosciences). Internalization of RBCs was assessed as previously described^2^. Briefly, *in vitro* generated RBCs were opsonized with anti-human RBC antibody (diluted 1:250; Rockland), stained with 2 µM CFSE (Thermo-Fisher) and then co-cultured for 18 h with primed Mo at a 1:2 (Mo to RBCs ratio) in cRPMI medium.

### LDH Cytotoxicity Assay and Propidium Iodide (PI) Permeability Assay

Primed BLaER1 Mo (50.000 cells in 150 μL of cRPMI medium with 2% FBS) were activated for the indicated time points. For LDH assay, cell culture supernatants were cleared of cells by spinning 96 well plates at 400 x g for 5 min. Supernatants were transferred to round bottom, non-treated 96 well plates and assayed for LDH release per the manufacturer’s protocol from the Sigma LDH cytotoxicity colorimetric assay kit. LDH release was expressed as LDH release (% vs Control) = [(unstimulated control − treated sample) * 100]/unstimulated control. For PI assay, cell pellet was resuspended in warmed plate reader media (HBSS + 20 mM HEPES + 10% FBS) supplemented with PI (5 µM). PI internalization was assessed by reading the fluorescence with an excitation wavelength of 530 nm and emission wavelength of 617 nm. PI incorporation was expressed as PI fluorescence (% vs Control) = [(unstimulated control − treated sample) * 100]/unstimulated control.

### Caspase-1 activity assay

Active caspase-1 was quantified by using the Caspase-1 Assay Kit Colorimetric (Abcam) according to the manufacturer’s instructions.

### Trim-Away

Purified human Mo were electroporated with mouse anti-AIM2 (10M5G5; Novus Biological), goat anti-MxA (R&D) or control IgG antibody. All antibodies used for electroporation were passed through Amicon Ultra-0.5 100 kDa centrifugal filter devices (Millipore) to remove traces of azide and replace buffer with PBS. Antibody electroporation was performed using the Neon Transfection System (Thermo Fisher). Mo were washed with PBS and resuspended in Buffer R (Thermo Fisher) at a concentration of 1x10^6^ cells/10 μL. For each electroporation reaction 10 µl of cells were mixed with 2 µg of antibody. The mixture was taken up into a 10 µl Neon Pipette Tip (Thermo Fisher) and electroporated using the following settings: 1400V, 20 ms, 2 pulses. Electroporated cells were transferred to cRPMI medium without antibiotics containing 10 ng/mL hrM-CSF (Stem Cell Technologies) and rested at 37°C for 1 h before stimulation. Rabbit anti-AIM2 (D5X7K; Cell Signaling) and Rabbit anti-MxA (Abcam) were use for Wester blot analysis of protein knockdown.

### Proteinase K (PK) protection assay of the 100k P fraction

PK protection assay was done as previously described with modifications^39^. In brief, the 100k P, obtained from 8×10^7^ BLaER1 cells upon differential centrifugation, was divided into three 30 µL fractions and treated as follows: without PK, with PK (15 µg/ml), and with PK (15 µg/ml) and 0.1% Triton X-100. The reactions were performed on ice for 20 min and stopped by adding 5× Lane Marker Reducing Sample Buffer (Thermo Fisher), supplemented with phenylmethylsulfonyl fluoride (PMSF; Thermo Fisher), and boiling for 5 min at 100°C. Samples were then subjected to SDS-PAGE followed by immunoblot analysis with the indicated antibodies.

### In vitro Proteoliposome (PtLp) translocation assay

The PtLp translocation assay was performed as previously described with some modifications^39^. Briefly, 1.25 mg of Bovine Liver Total Lipid Extract (Avanti) in chloroform was evaporated by a stream of nitrogen gas over and further dried in 37°C incubator for 1 h. Dried lipids were suspended in 400 μL of HEPES-KAc buffer contain 20mM HEPES (pH 7.2) and 150mM potassium acetate (HEPES-KAc buffer). The lipids were repeatedly frozen in liquid nitrogen and thawed in 42°C water bath for 10 times. Triton X-100 (TX100) was then added into the lipid solution to a final concentration of 0.05% and rotated in 4°C for 30 min. Recombinant MxA (5 μg; Novus Biological) was then added into the lipid/TX100 solution and incubated for another 1 h with rotation. Each 400 µL solution was incubated with 5 mg Biobeads SM2 (Bio-rad) equilibrated with the HEPES-KAc buffer at 4°C. Beads were replaced each hour and repeated for 5 times (10 mg beads in the four time and incubated overnight). After a 1,500 g centrifugation to remove the Biobeads, the PtLp solution was repeatedly frozen in liquid nitrogen and thawed in 42°C water bath for 5 times. In order to remove the free proteins, the PtLp solution was centrifuged at 100,000 g for 1 h and the supernatant discarded. For the *in vitro* mIL-1β translocation, the PtLp pellet was resuspend in 90 µL of HEPES-KAc buffer with 1 µg mIL-1β (Invivogen) and incubated for 1 h in 30°C. The solution was aliquoted into 3 fractions (30 µL each). The first fraction was a control, the second and third fractions were digested by PK (10 µg/ml) without or with 0.5% TX100 for 20 min on ice. The reactions were stopped by 1 mM PMSF and incubated for 10 min on ice. Then 5× Lane Marker Reducing Sample Buffer (Thermo Fisher) was added, and the samples were heated at 100°C for 10 min followed by immunoblot analysis. To assess the stable incorporation of MxA within the PtLp a membrane flotation procedure was performed. Briefly, to 350 µL of PtLp solution (incubated for 5 min at room temperature with or without 2 M urea), 350 µL 50% OptiPrep (diluted in HEPES-KAc buffer) was added. The mixture was overlaid with 560 µL 20% OptiPrep and 105 µL HEPES-KAc buffer, centrifuged at 100,000 g for 2 h. Five fractions were collected and subjected to immunoblot analysis.

### Release of encapsulated mIL-1β or LDH from the lumen of liposomes (Lp)

The release of encapsulated mIL-1β from the lumen of Lp was performed as previously described with some modifications^33^. 2.5 mg of Bovine Liver Total Lipid Extract (Avanti) in chloroform was evaporated by a stream of nitrogen gas and further dried in 37°C incubator for 1 h. Dried lipid film was resuspended in 500 μL of buffer containing 25 mM Tris-HCl (pH 8.0) - 150 mM NaCl (Lp buffer) and 2 µg of recombinant human mIL-1β (Invivogen) or 10 µg of LDH (Sigma). To increase reconstitution of dried lipid film with recombinant proteins and buffer, the tube containing lipid, buffer, and recombinant protein was covered and incubated at 37°C for 15 min. The combined solution was then vortexed continuously for 5 min at room temperature to encapsulate recombinant proteins into large Lp. The Lp solution was spun at 100,000 x g for 15 min at 4°C to pellet Lp containing the recombinant protein. Supernatant containing the free protein was removed. The Lp pellet was gently washed 3 times with Lp buffer. After the final wash, the Lp pellet was resuspended in a final volume of 1200 µL of Lp buffer for protein release experiments. 170 µL of the Lp solution were aliquoted into 6 tubes. Recombinant MxA (5 μg; Novus Biological) or Lp buffer were then added to Lp solution aliquots for indicated time points. The solution was then spun at 100,000 x g for 30 min and the supernatants treated with 5× Lane Marker Reducing Sample Buffer (Thermo Fisher) for subsequent Western blotting. One tube corresponding to the maximal pellet signal and minimal supernatant signal was spun at 100,000 x g for 30 min immediately after MxA was added to the Lp solution. The corresponding pellet and supernatant were then used as a positive and negative control respectively. For LDH experiments, resuspended pellet was lysed with 10X lysis buffer from the LDH colorimetric quantification kit (Sigma) and liposome buffer to make a final volume equivalent to harvested supernatants. 50 µL of lysed pellet and supernatants were incubated with 50 µL of colormetric LDH substrate for 30 min to assay LDH activity. Absorbance values were normalized to the lysed pellet (set to 100% activity).

### RNA preparation and sequencing library preparation

Total RNA was isolated from cell lysates using the RNeasy Micro Kit (QIAGEN), including on-column deoxyribonuclease digestion, and was analyzed for quality using the RNA 6000 Pico Kit (Agilent Technologies). Poly(A)-enriched next-generation sequencing library construction was performed using the KAPA mRNA Hyper Prep Kit (KAPA Biosystems) with 100 ng of input total RNA and 15 amplification cycles according to the manufacturer’s protocol. The quality and quantity of the individual libraries was assessed using the High Sensitivity DNA Kit (Agilent). The libraries were equimolar pooled, and an additional 1X bead clean-up was used to remove adaptor dimer. The pooled libraries were quantitated using the KAPA Library Quantification Kit, Universal (KAPA Biosystems) and sequenced on an Illumina NextSeq 500 with paired-end 75-base-pair (bp) read lengths.

### RNA sequencing data processing and analysis

Quality control of raw reads was performed with FastQC. Reads were aligned to the reference human genome (GRCh38) using HISAT2 after quality and adaptor trimming by Cutadapt. After sorting binary alignment map files by name using samtools, the HTSeq-count program was used to quantify total numbers of read counts mapped to the genome. The RNA sequencing data analysis was performed in the R programming language. The DESeq2 R package was used for size factor and dispersion estimation calculation and differential gene expression analysis.

### Quantification and statistical analysis

All data are presented as means ± SEM from n=biological replicates (for experiments involving BLaER1 cells) or n=independent donors (for experiments involving human primary cells). Statistical analyses were performed using GraphPad Prism (version 9, GraphPad Software). Comparison of two parameters was performed using two-tailed unpaired Student t test. Multiparameter analyses of cell-based studies were performed with one-way repeated measures analysis of variance (ANOVA) with Greenhouse-Geisser correction. Statistical significance was defined as n.s. = not significant.

## ACKNOWLEDGMENTS

We are grateful to our SLE and JDM patients, their families, the healthy individuals who participated in our study and the members of the Pediatric Rheumatology Clinics at Texas Scottish Rite Hospital for Children and the Children’s Medical Center in Dallas (TX). We thank Dr. L. Cohen-Gould and Dr. S. Mukherjee (WCM− New York, NY) for help with microscopy studies, Thomas Miller (Drukier Institute for Children’s health − WCM, New York, NY) for help with flow cytometry and cell sorting, the Genomics Core at the Baylor Scott and White Research Institute (Dallas, TX) for RNA-seq and the Genomics Core at the Jackson Laboratory for Genomic Medicine (Farmington, CT) for scRNA-seq.

## Funding

This work was supported by NIAMS CORT P50AR070594 Center for Lupus Research (to V.P. and J.F.B.), NIAID NIH U19 AI082715 (to V.P.), Lupus Research Alliance − LRA 704798 (to S.C.) and funds from the Drukier Institute for Children’s Health at Weill Cornell Medicine.

## Author contributions

S.C., P.B., J. R-A, Z.W. and R.M. performed the experiments. J.B., L.W., P.S., C.S., L.N., K.S., J.F., and T.W. coordinated the collection of human samples and provided clinical information. U.B., D.N-B. and J.G. analyzed RNA-seq and scRNA-seq data and provided recommendations for statistical analysis. S.C. and V.P. conceptualized the study, designed experiments and interpreted data. S.C, V.P. and J.F.B. supervised the study and wrote the manuscript.

## Competing interests

V.P. has received consulting honoraria from Sanofi, Astra-Zeneca and Moderna and is the recipient of a research grant from Sanofi and a contract from Astra Zeneca. J.F.B. is an employee of Immunai Inc. V.P. and J.F.B. receive royalties for use of Canakinumab in sJIA.

---

**Fig. S1.**
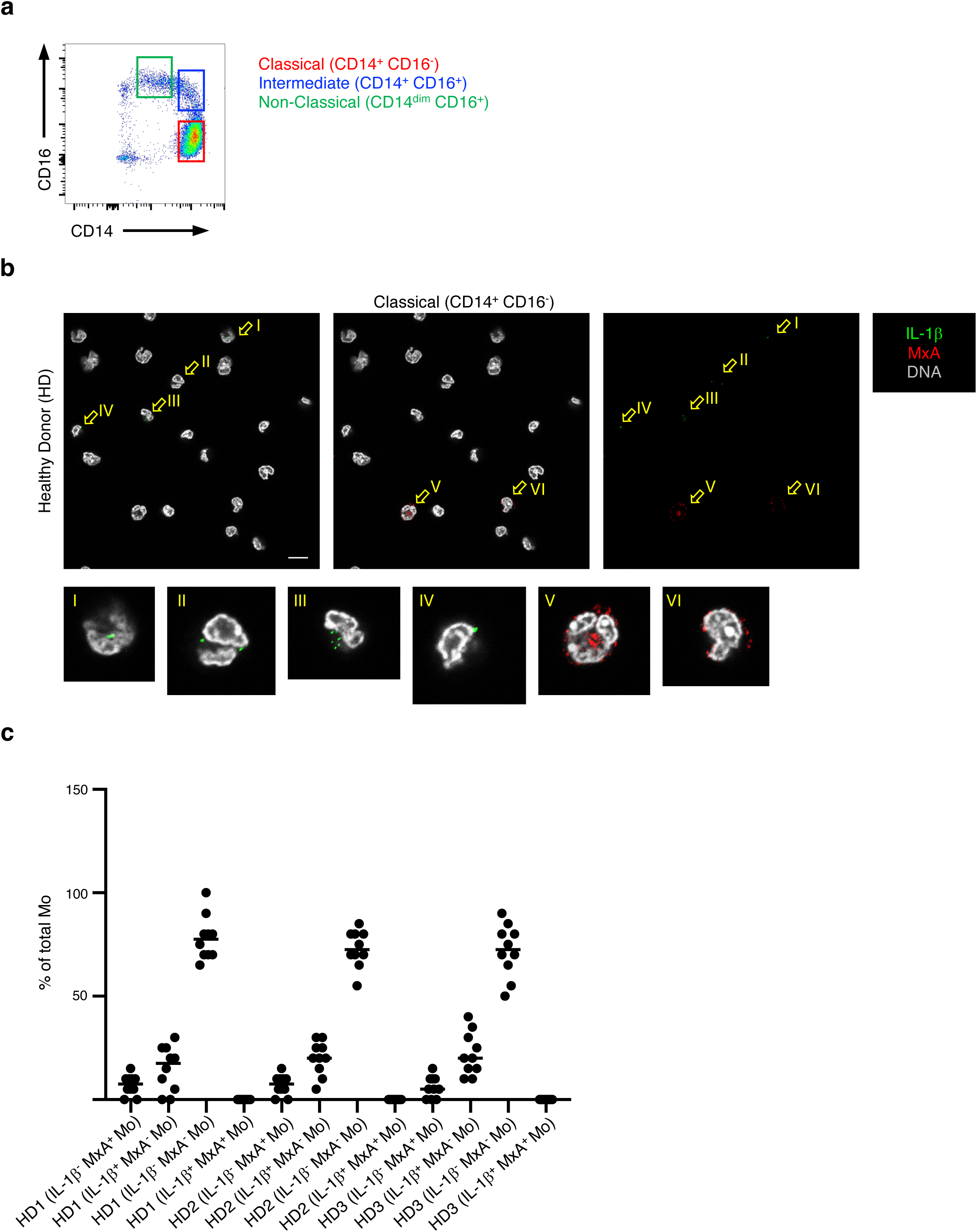
**a,** Sorting strategy used to purify blood Mo subsets. **b,** Confocal images of classical Mo isolated from healthy donors (HD). Scale bar: 20 µm. **c,** Percentages of IL-1β^−^ MxA^+^, IL-1β^+^ MxA^-^, IL-1β MxA^-^ and IL-1β^+^ MxA^+^ Mo from three HD. Each dot represents the count from different micrscopy fields from a study participant sample that was processed, stained and analyzed independently.

**Fig. S2.**
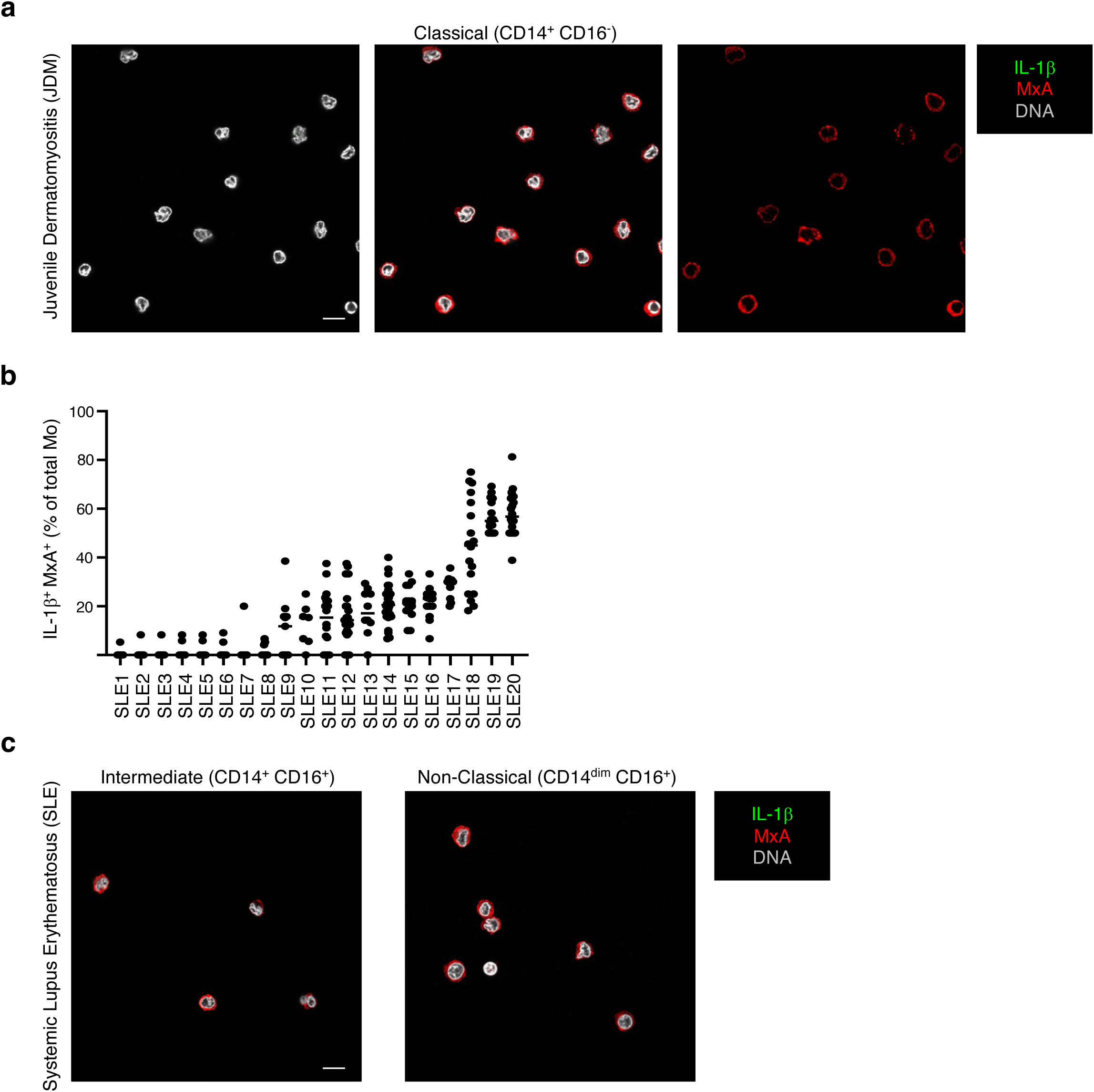
**a,** Confocal images of classical Mo isolated from JDM patients. Scale bar: 20 µm. **b,** Percentages of IL-1β^+^ MxA^+^ Mo from the twenty SLE patients show in Fig. 1a. Each dot represents the count from different micrscopy fields from a study participant sample that was processed, stained and analyzed independently. **c,** Confocal images of intermediate and non-classical Mo isolated from SLE patients. Scale bar: 20 µm.

**Fig. S3.**
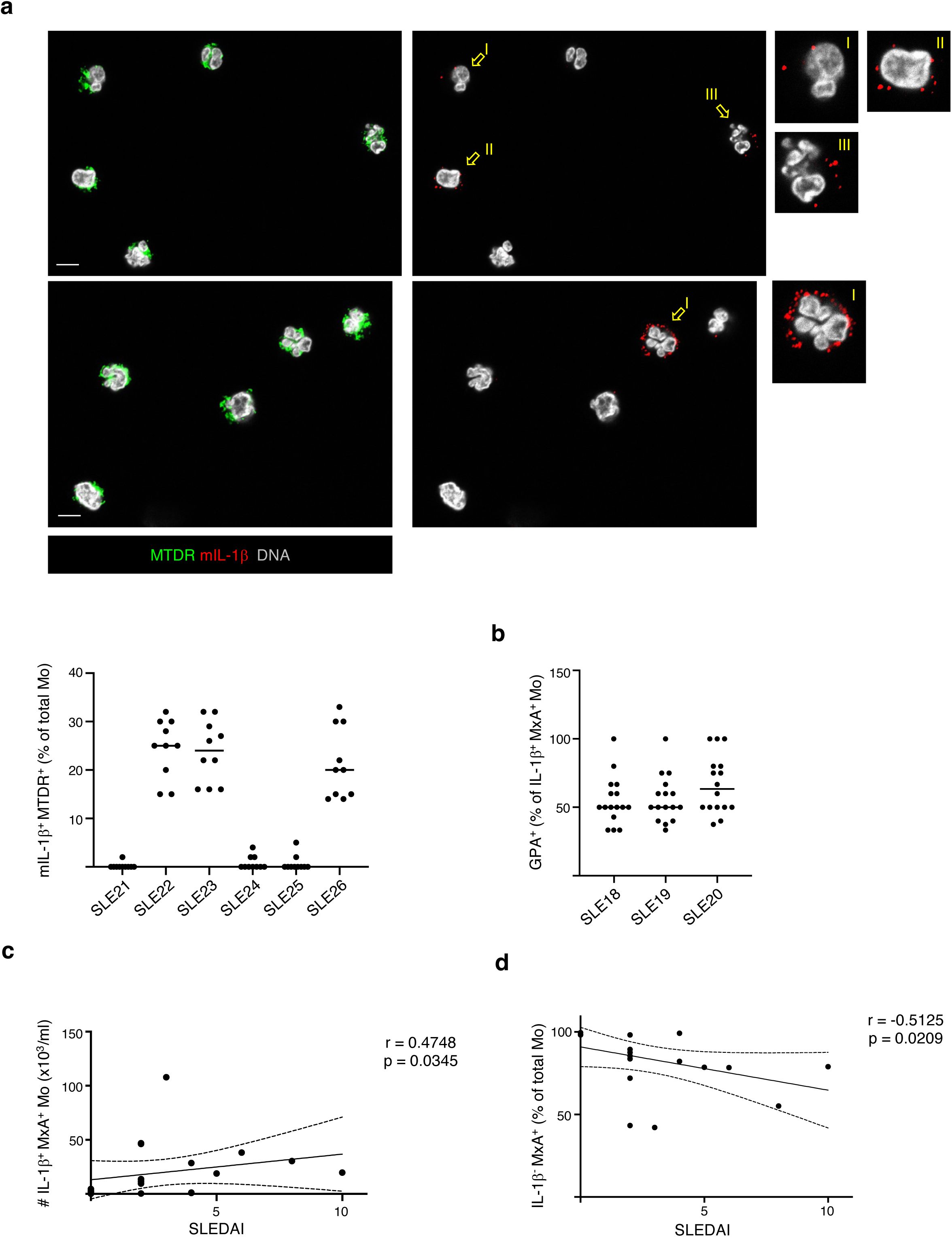
**a,** Representative confocal images of extracellular mIL-1β from classical Mo sorted from SLE patient. Slides were coated with anti-mIL-1β capture antibody and mitochondria were stained with MitoTracker Deep Red (MTDR). Scale bar: 20 µm. The percentages of mlL-1β^+^ MTDR^+^ Mo from six SLE is also shown. Each dot represents the count from different micrscopy fields from a study participant sample that was processed, stained and analyzed independently. **b,** Percentages of GPA^+^ Mo, within IL-1β^+^ MxA^+^ Mo, from three SLE patients. Each dot represents the count from different micrscopy fields from a study participant sample that was processed, stained and analyzed independently. **c,** Two-tailed Pearson’s correlation between SLEDAl and the absolute number (#) of IL-1β^+^ MxA^+^ Mo (n=20). Dashed lines represent 95% confidence intervals. **d,** Two-tailed Pearson’s correlation between SLEDAl and the percentage of lL-1β^-^ MxA^+^ Mo (n=20). Dashed lines represent 95% confidence intervals.

**Fig. S4.**
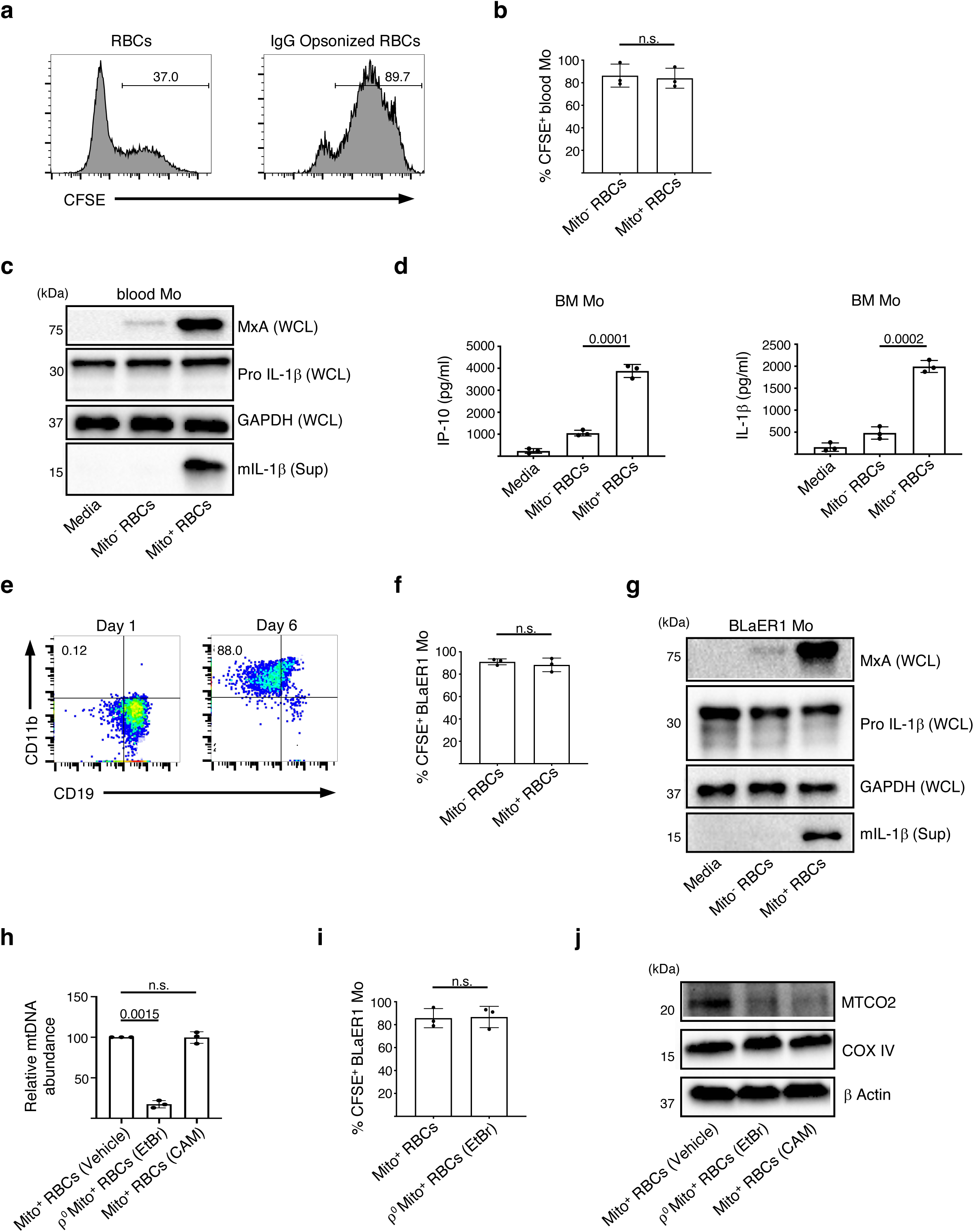
**a,** Phagocytosis of CFSE-labeled RBCs in the presence or absence of IgG opsonizing antibodies. **b,** Quantification of phagocytosis of CFSE-labeled Mito^-^ or Mito^+^ RBCs by blood Mo. (n=3). **c,** Immunoblot analysis of whole cell lysate (WCL) and supernatants (Sup) from blood Mo that were treated as described. One representative of two experiments. **d,** Levels of IP-10 and IL-1β in the supernatants of bone marrow (BM) Mo cultured with media, Mito^-^ or Mito^+^ opsonized RBCs. (n=3). **e,** Phenotype of BLaER1 cells before (Day 1) and after (Day 6) differentiation into BLaER1 Mo. **f,** Quantification of phagocytosis of CFSE-labeled Mito^-^ or Mito^+^ RBCs by BLaER1 Mo. (n=3). **g,** Immunoblot analysis of WCL and Sup from BLaER1 Mo that were treated as described. One representative of two. **h,** Relative mtDNA abundance in Mito^+^ RBCs generated with vehicle, EtBr (ρ^0^) or CAM. (n=3). **i,** Quantification of phagocytosis of CFSE-labeled Mito^+^ RBCs RBCs generated with vehicle or EtBr (ρ^0^). (n=3). **j,** Western blot analysis of Mito^+^ RBCs generated with vehicle, EtBr (ρ^0^) or CAM. One representative of two.

**Fig. S5.**
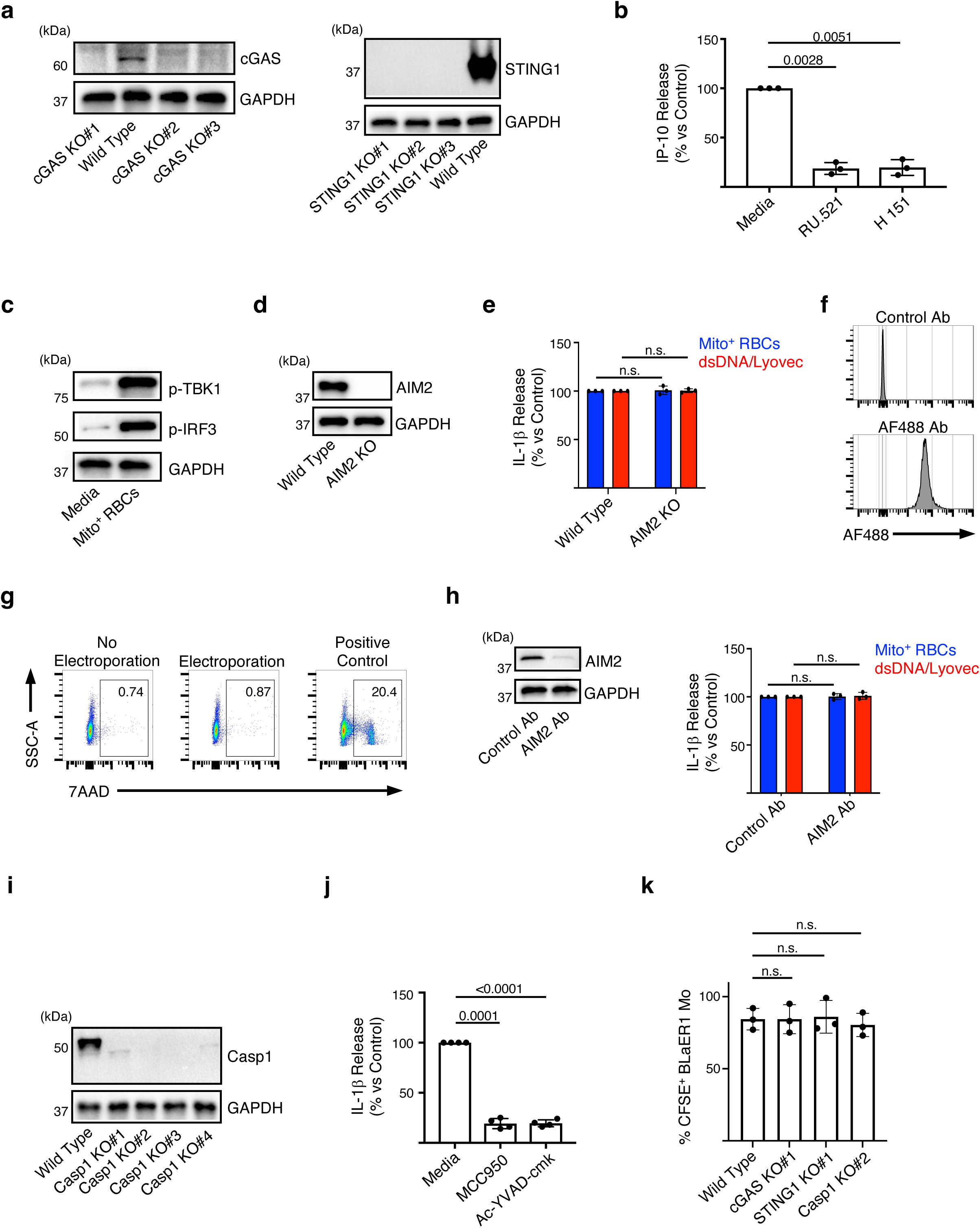
**a,** Western blot analysis confirming CRISPR/Cas9-mediated KO of cGAS and STING1. **b,** Normalized IP-10 levels in the supernatants of Mito^+^ RBC-activated BLaER1 Mo in the presence of the cGAS inhibitor RU.521 or the STING inhibitor H151. (n=3). **c,** Western blot analysis of phosphorylated TBK1 and IRF3 in BLaER1 Mo cultured with media or Mito^+^ RBCs. **d,** Western blot analysis confirming CRISPR/Cas9-mediated KO of AIM2. **e,** Normalized IL-1β levels in the supernatants of Mito^+^ RBCs-activated wild type or AIM2 KO BLaER1 Mo. (n=3). **f,** Flow cytometry analysis showing the internalization of unlabeled (control) or AlexaFluor 488 (AF488)-labeld IgG upon electroporation. **g,** Percentages of cell death measured by 7AAD incorporation. **h,** Western blot analysis confirming Trim-Away knockdown of AIM2 (left). Normalized IL-1β levels in the supernatants of activated BLaER1 Mo upon Trim-Away knockdown of AIM2 (right). (n=3). **i,** Western blot analysis confirming CRISPR/Cas9-mediated KO of Caspase-1. **j,** Normalized IL-1β levels in the supernatants of Mito^+^ RBCs-activated BLaER1 Mo in the presence of the NLRP3 inhibitor MCC950 or the caspase-1 inhibitor Ac-YVAD-cmk. (n=4). **k,** Phagocytosis of CFSE-labeled RBCs in wild type, cGAS KO, STING1 KO or caspase-1 KO BLaER1 Mo. (n=3).

**Fig. S6.**
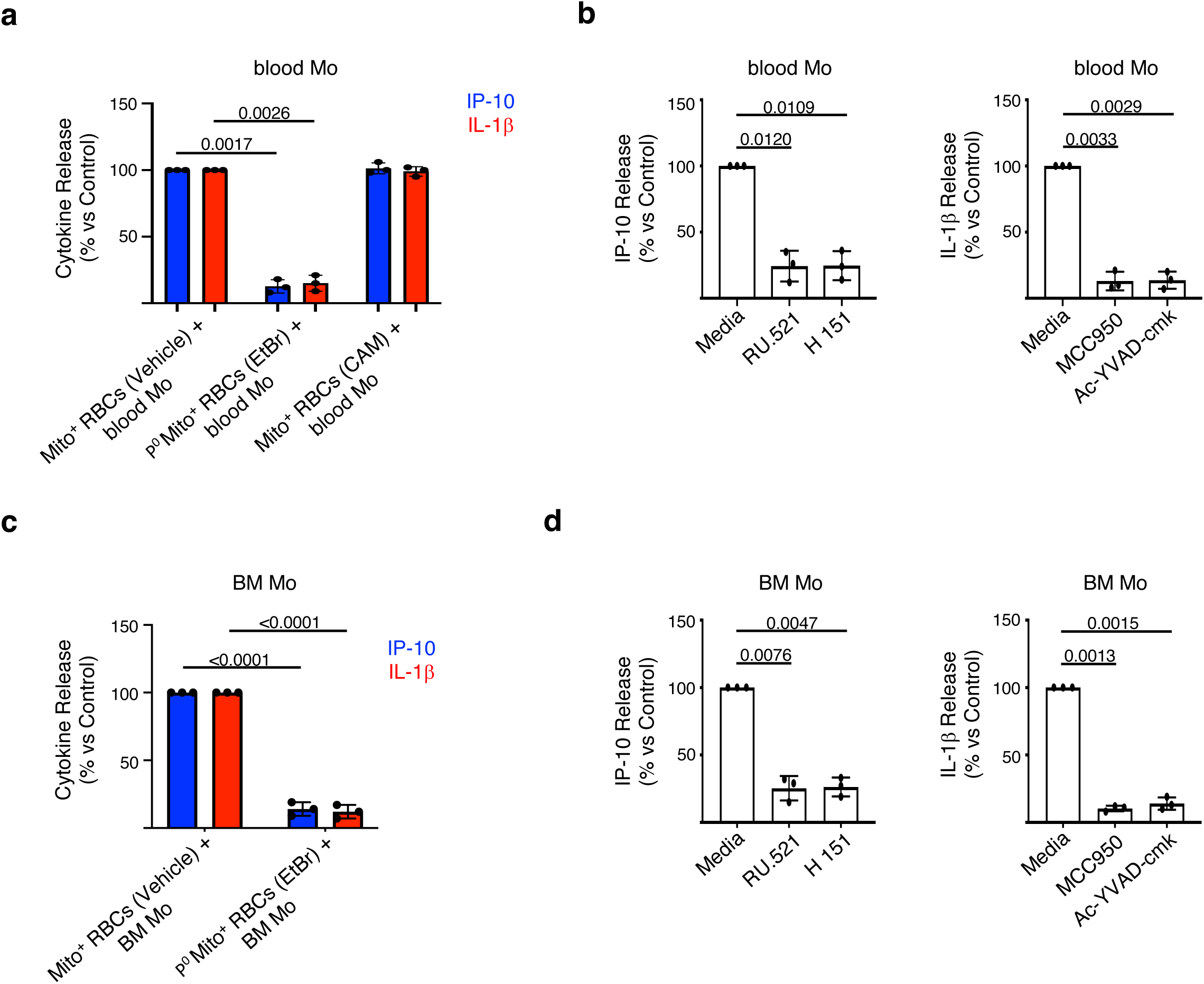
**a,** Normalized cytokine levels in the supernatants of blood Mo activated with Mito^+^ RBCs generated in the presence of vehicle, EtBr (ρ^0^) or CAM. **b,** Normalized cytokine levels in the supernatants of Mito^+^ RBC-activated blood Mo in the presence of RU.521, H151, MCC950 or Ac-YVAD-cmk. (n=3). **c,** Normalized cytokine levels in the supernatants of BM Mo activated with Mito^+^ RBCs generated in the presence of vehicle, EtBr (ρ^0^) or CAM. **d,** Normalized cytokine levels in the supernatants of Mito^+^ RBCs-activated BM Mo in the presence of RU.521, H151, MCC950 or Ac-YVAD-cmk. (n=3).

**Fig. S7.**
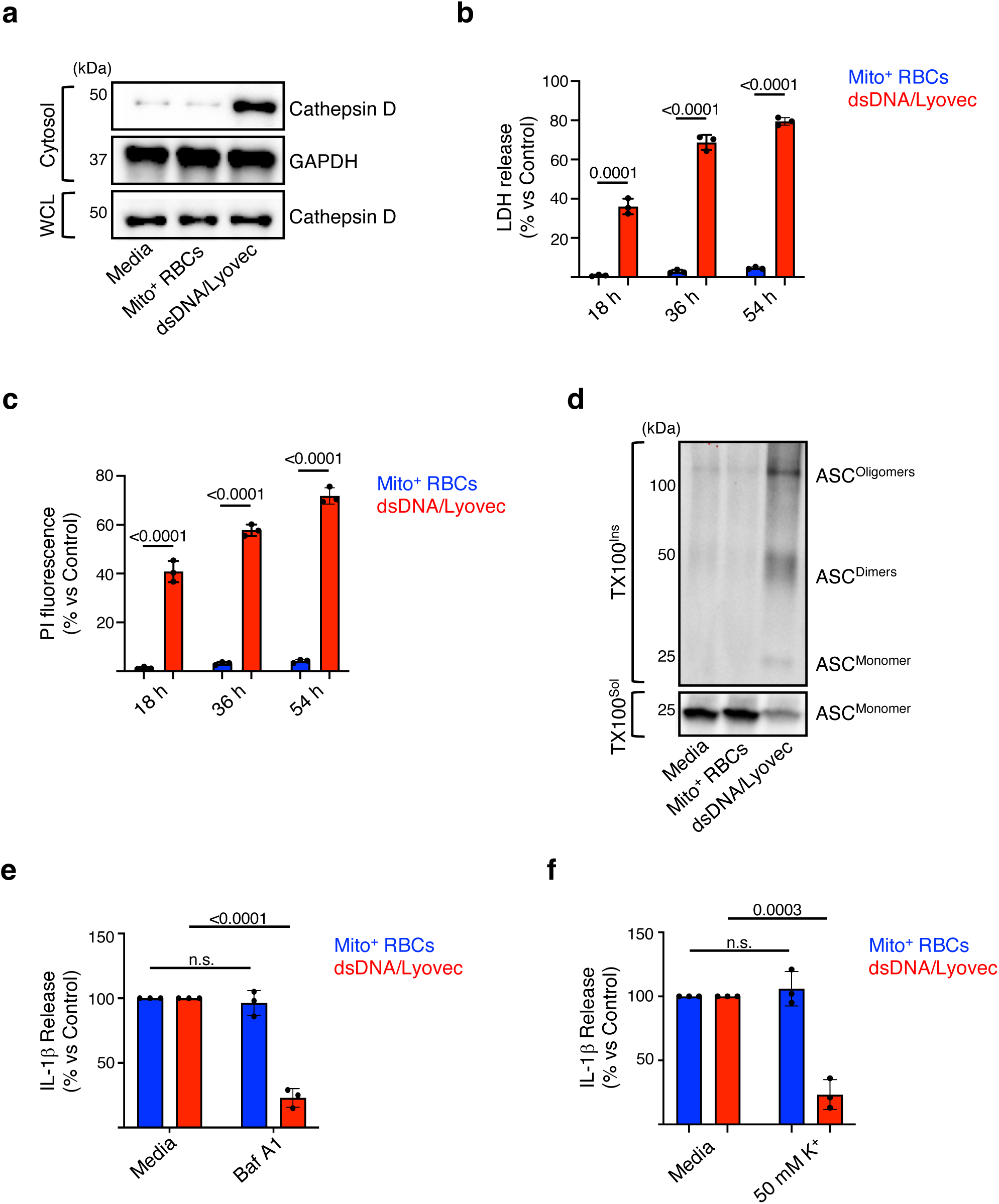
**a,** Western blot analysis of cytosolic fraction obtained from media, Mito^+^ RBCs- or dsDNA/Lyovec-activated BLaER1 Mo. WCL: whole cell lysate. Quantification of lactate dehydrogenase (LDH) release **(b)** or propidium iodide (PI) internalization **(c)** in BLaER1 Mo activated Mito^+^ RBCs or dsDNA/Lyovec over time. Data are normalized to media-treated cells. (n=3). **d,** Western blot analysis of ASC oligomers (pyroptosome) formation in BLaER1 Mo activated Mito^+^ RBCs or dsDNA/Lyovec for 18 h. Triton X-100 soluble (TX100^Sol^) and insoluble (TX100^Ins^) fractions were immunoblotted with ASC antibody. One representative of two experiments. **e,** Normalized IL-1β levels in the supernatants of Mito^+^ RBC- or dsDNA/Lyovec-activated BLaER1 Mo in the presence of the lysosmal acidification inhibitor Bafilomycin A1 (BAf A1). (n=3). **f,** Normalized IL-1β levels in the supernatants of Mito^+^ RBC- or dsDNA/Lyovec-activated BLaER1 Mo in the presence of extracellular K^+^. (n=3).

**Fig. S8.**
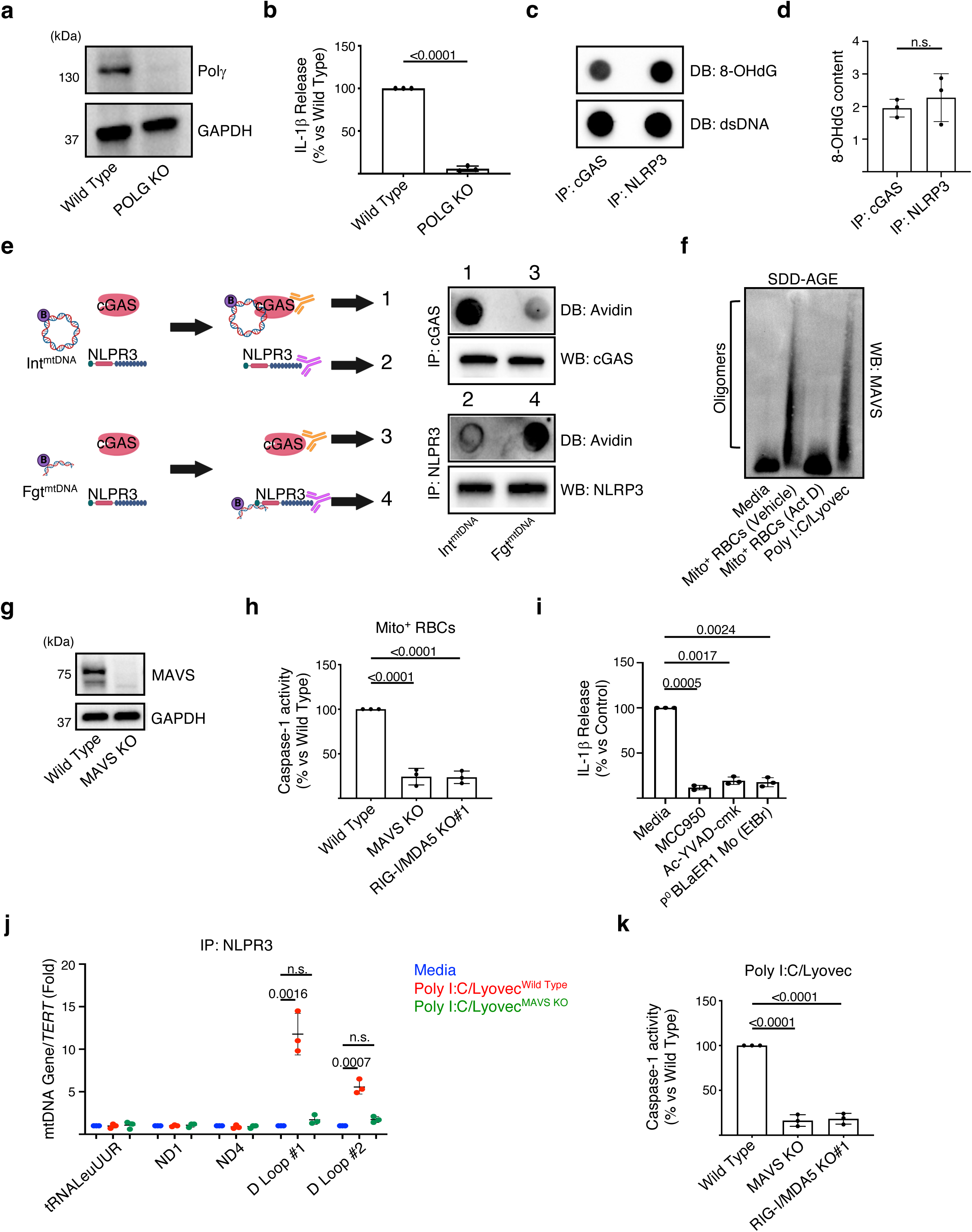
**a,** Western blot analysis confirming CRISPR/Cas9-mediated KO of mitochondrial DNA polymerase gamma (POLG; Polγ). **b,** Normalized IL-1β levels in the supernatants of Mito^+^ RBC-activated wild type or POLG KO BLaER1 Mo. (n=3). **c,** DB analysis of cGAS or NLRP3 immunocomplexes isolated from Mito^+^ RBCs-activated BLaER1 Mo. As a loading control, dsDNA dot blot (DB) was performed. **d,** 8-OHdG quantification by ELISA in cGAS or NLRP3 immunocomplexes isolated from Mito^+^ RBC-activated BLaER1 Mo. (n=3). **e,** Experimental scheme (left) and DB analysis (right) of IP cGAS or NLRP3 upon incubation with biotinylated intact (Int) or fragmented (Fgt) mtDNA. As a loading control, cGAS or NLRP3 immunoblot was performed. **f,** SDD-AGE of MAVS oligomerization in BLaER1 Mo in response to Mito^+^ RBCs (generated with or without Act D) or to transfection with Poly I:C. One representative of three. **g,** Western blot analysis confirming CRISPR/Cas9-mediated KO of MAVS. **h,** Relative caspase-1 activity in the lysate of Mito^+^ RBC-activated wild type, MAVS or RIG-I/MDA5 KOBLaER1 Mo. (n=3). **i,** Normalized IL-1β levels in the supernatants of Poly I:C/Lyovec-activated BLaER1 Mo in the presence of MCC950 or Ac-YVAD-cmk or that were generated in the presence of EtBr. (n=3). **j,** Relative levels of mtDNA-encoded genes in NLRP3 immunocomplexes isolated from Poly I:C/Lyovec-activated wild type or MAVS KO BLaER1 Mo. (n=3). **k,** Relative caspase-1 activity in the lysate of Poly I:C/Lyovec-activated wild type, MAVS or RIG-I/MDA5 KO BLaER1 Mo. (n=3).

**Fig. S9.**
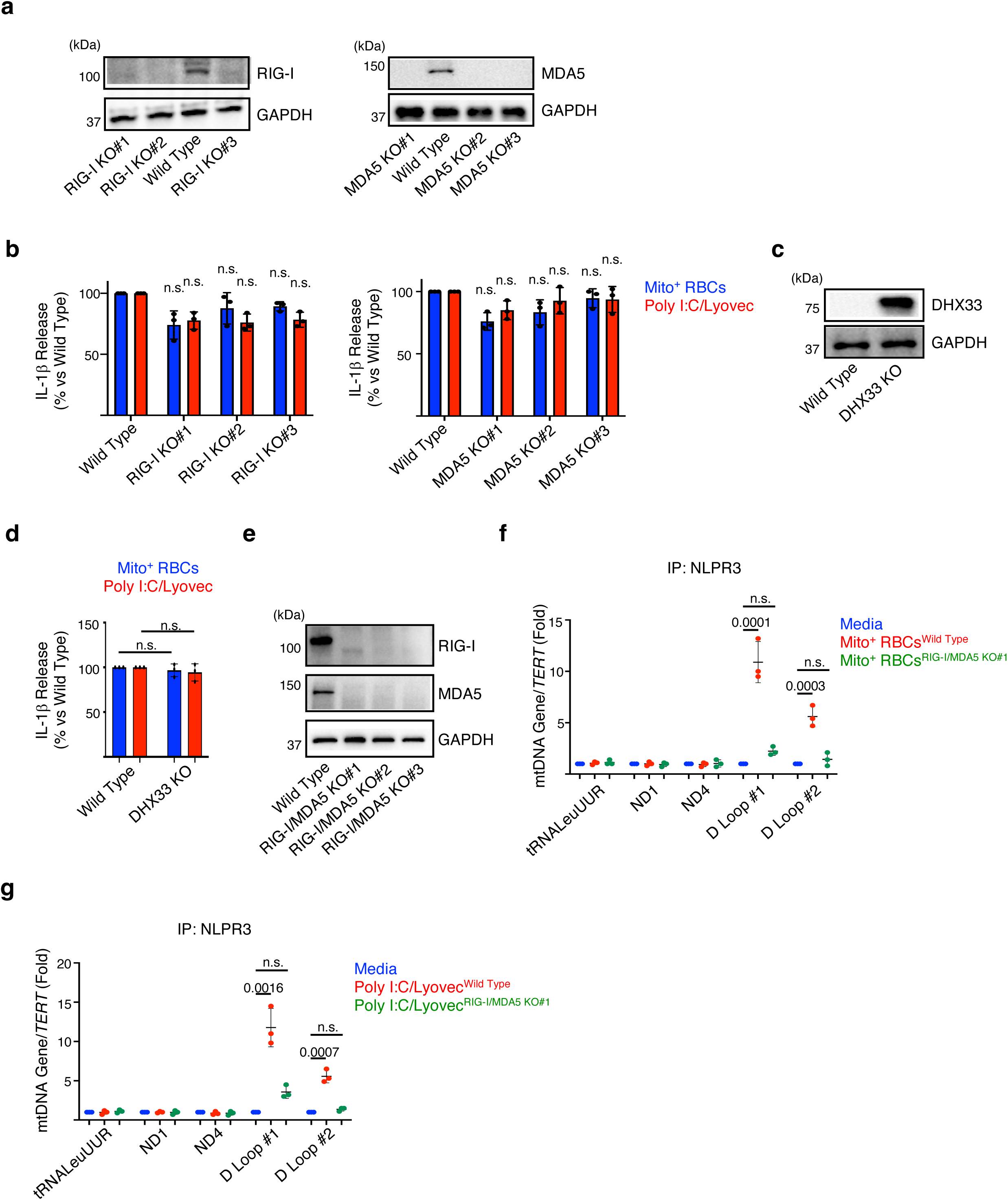
**a,** Western blot analysis confirming CRISPR/Cas9-mediated KO of RIG-I or MDA5. **b,** Normalized IL-1β levels in the supernatants of Mito^+^ RBCs or Poly I:C/Lyovec-activated wild type, RIG-I KO or MDA5 KO BLaER1 Mo. (n=3). **c,** Western blot analysis confirming CRISPR/Cas9-mediated KO of DHX33. **d,** Normalized IL-1β levels in the supernatants of Mito^+^ RBCs or Poly I:C/Lyovec-activated wild type or DHX33 KO BLaER1 Mo. (n=3). **e,** Western blot analysis confirming CRISPR/Cas9-mediated KO of both RIG-I and MDA5. Relative levels of mtDNA-encoded genes in NLRP3 immunocomplexes isolated from Mito^+^ RBCs **(f)** or Poly I:C/Lyovec **(g)** activated wild type or RIG-I/MDA5 KO BLaER1 Mo. (n=3).

**Fig. S10.**
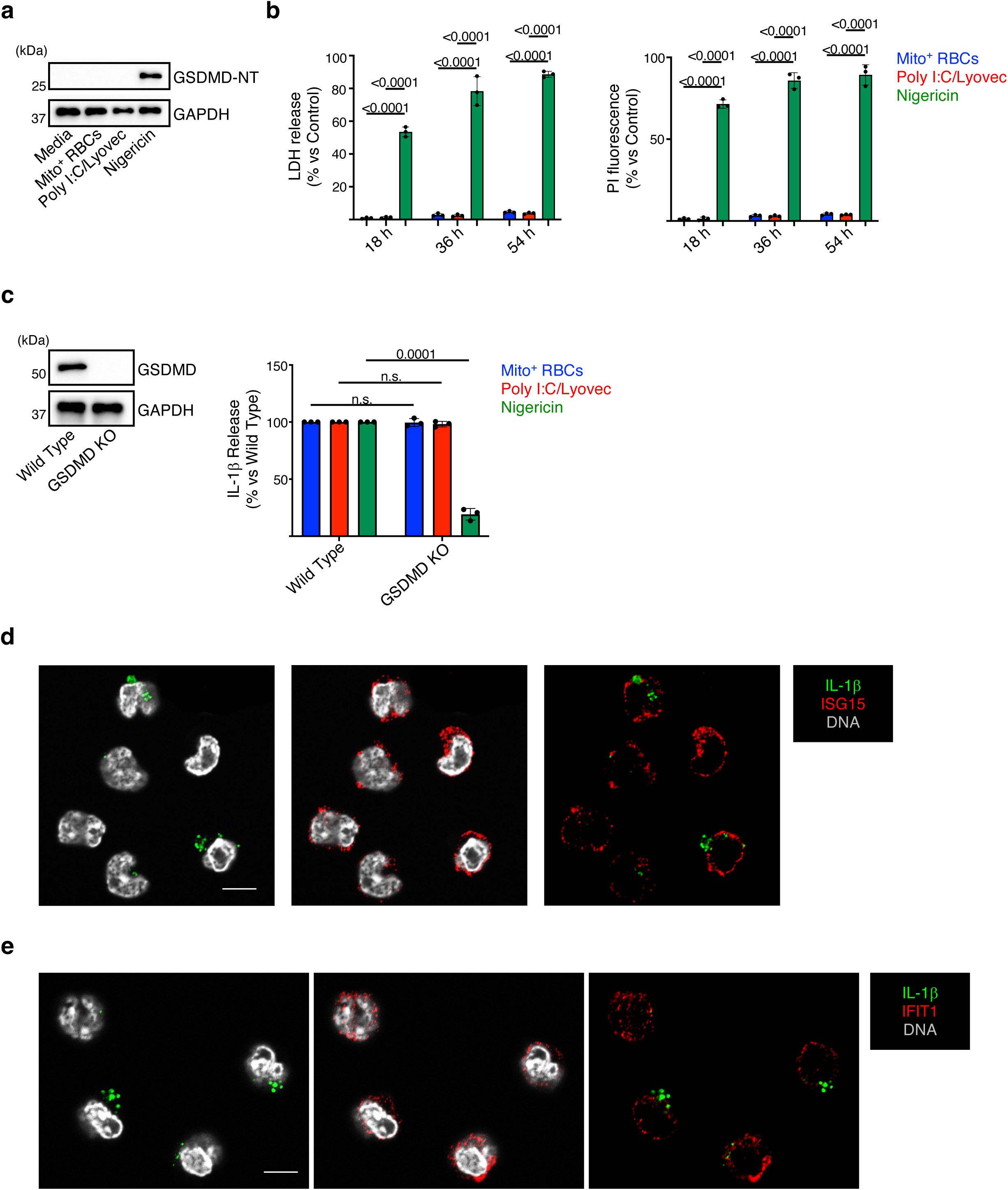
**a,** Western blot analysis of gasdermin-D N-terminal fragment (GSDMD-NT) in total cell lysate isolated from Mito^+^ RBCs, Poly I:C/Lyovec and Nigericin-activated BLaER1 Mo. **b**, Quantification of lactate dehydrogenase (LDH) release or propidium iodide (PI) internalization in BLaER1 Mo activated Mito^+^ RBCs, Poly I:C/Lyovec or Nigericin over time. Data are normalized to media-treated cells. (n=3). **c,** Western blot analysis confirming CRISPR/Cas9-mediated KO of GSDMD (left). Normalized IL-1β levels in the supernatants from Mito^+^ RBCs, Poly I:C/Lyovec and Nigericin-activated wild type or GSDMD KO BLaER1 Mo (right). (n=3). Confocal images of classical Mo isolated from SLE patients and stained for IL-1β and ISG15 **(d)** or IFIT1 **(e).** Scale bar: 15 µm.

**Fig. S11.**
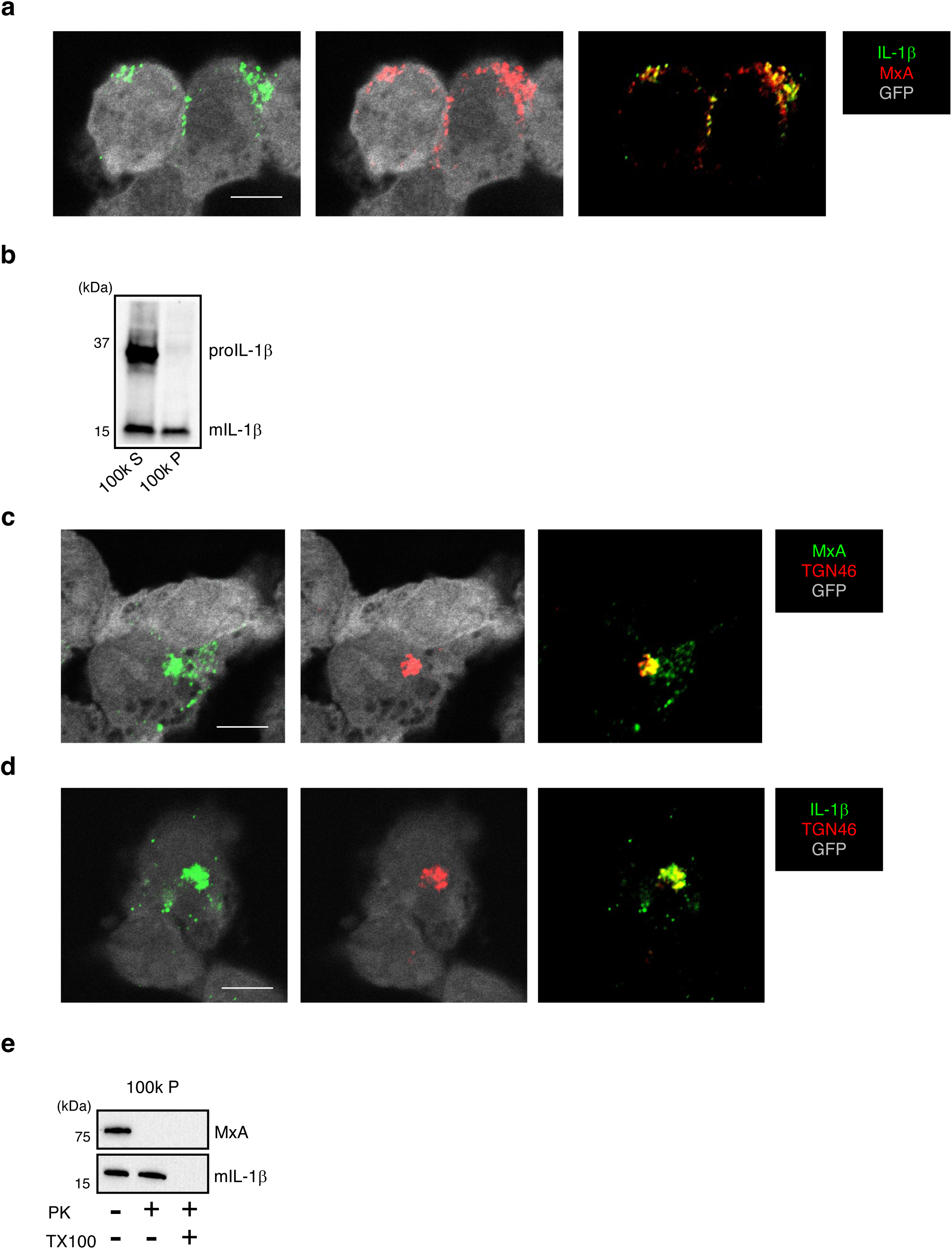
**a,** Confocal images of Poly I:C/Lyovec-activated BLaER1 Mo stained for IL-1β and MxA. Scale bar: 5 µm. **b,** Western blot analysis, with anti-IL-1β antibody (MAB201) of the microsomal pellet (100k P) and postmicrosomal supernatant (100k S) isolated from Poly I:C/Lyovec-activated BLaER1 Mo. One representative of two. Confocal images of Poly I:C/Lyovec-activated BLaER1 Mo stained for MxA and TGN46 **(c)** or IL-1β and TGN46 **(d)**. Scale bar: 5 µm. **e,** Proteinase K (PK) protection assay of the membranes fraction (100k P) isolated from Poly I:C/Lyovec-activated BLaER1 Mo. TX100: Triton X-100.

**Fig. S12.**
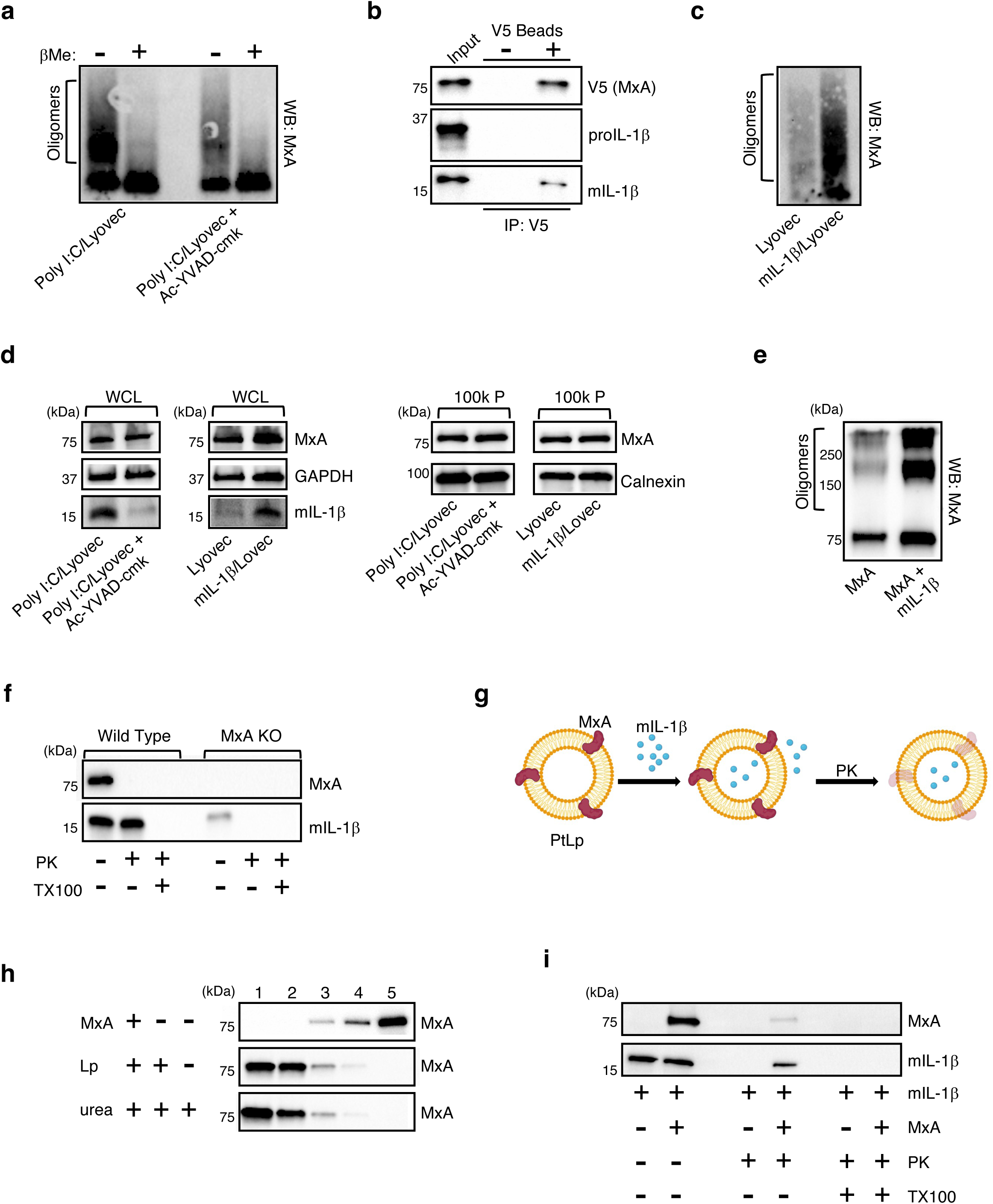
**a,** SDD-AGE of MxA oligomerization in the membranes fraction (100k P) isolated from Poly I:C/Lyovec-activated BLaER1 Mo with or without Ac-YVAD-cmk. βMe: βmercaptoethanol. One representative of two. **b,** co-IP using the lysate from MxA-V5 expressing, Poly I:C/Lyovec-activated, BLaER1 Mo with anti-V5 (+) or control (-) agarose beads. One representative of two. **c,** SDD-AGE of MxA oligomerization in the membranes fraction (100k P) isolated from INFα2β-primed BLaER1 Mo transfected with mIL-1β. One representative of two. **d,** Western blot analysis of the whole cell lysate (WCL; left) or of the membranes fraction (100k P; right) isolated from BLaER1 Mo treated as described. **e,** SDS-PAGE of *in vitro* MxA oligomerization in the presence of recombinant mIL-1β. One representative of three. **f,** PK protection assay of the membranes fraction isolated from Poly I:C/Lyovec-activated wild type or MxA KO BLaER1 Mo. TX100: Triton X-100. **g,** Experimental scheme of the *in vitro* proteoliposomes (PtLp)-based transport assay. **h,** Western blot analysis of a membrane floation assay performed with MxA alone or MxA PtLp that were treated with or without 2M urea. Lp: liposomes. **i,** Western blot analysis of the *in vitro* PtLp-based transport assay. In brief, MxA PtLp were incubated with recombinant human mIL-1β. PK digestion was then performed, with or without TX100, to determine the amount of mIL-1β that were incorporated within the PtLp lumen. One representative of two.

**Fig. S13.**
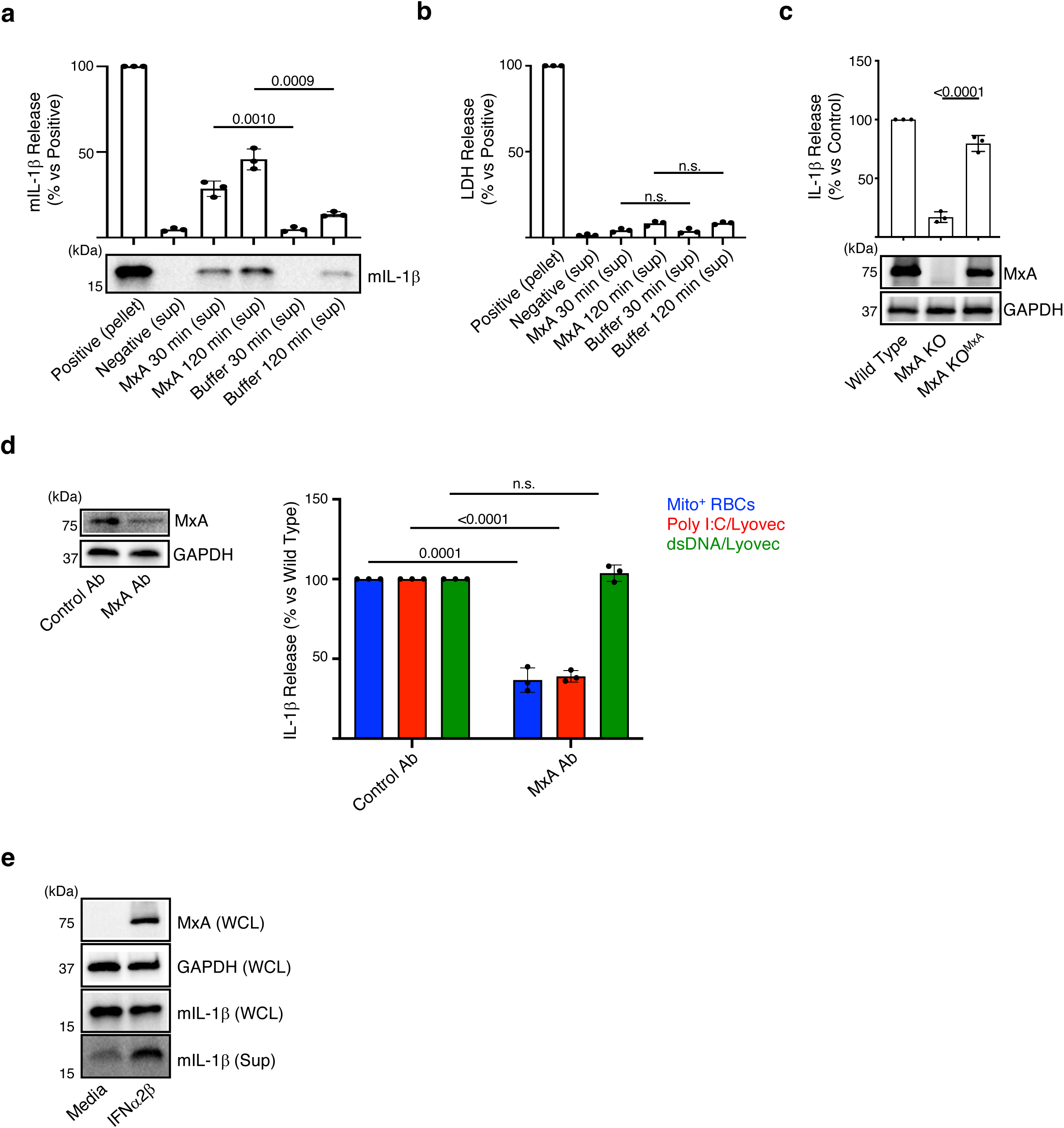
**a,** Immunoblot analysis of mIL-1β present within liposomes (pellet), or supernatants (sup) after ultracentrifugation of liposomes that were treated with buffer or recombinant human MxA for 0 (control), 30 and 120 min. Densitometry quantification of Western blot band density for mIL-1β release from liposomes is also shown. (n=3). **b,** Quantification of LDH enzymatic activity release from Lp loaded with LDH and treated with buffer or MxA for indicated times. (n=3). **c,** Normalized IL-1β levels in the supernatants from Mito^+^ RBCs-activated BLaER1 Mo with the indicated genotype. (n=3). Immunoblot analysis of the total cell lysate is also shown. **d,** Western blot analysis confirming Trim-Away knockdown of MxA (left). Normalized IL-1β levels in the supernatants of activated BLaER1 Mo upon Trim-Away knockdown of MxA (right). (n=3). **e,** Western blot analysis of whole cell lysate (WCL) and supernatants (Sup) isolated from BLaER1 Mo overexpressing mIL-1β that were treated with or without recombinant Type I IFN (IFNα2β). One representative of two.

**Fig. S14.**
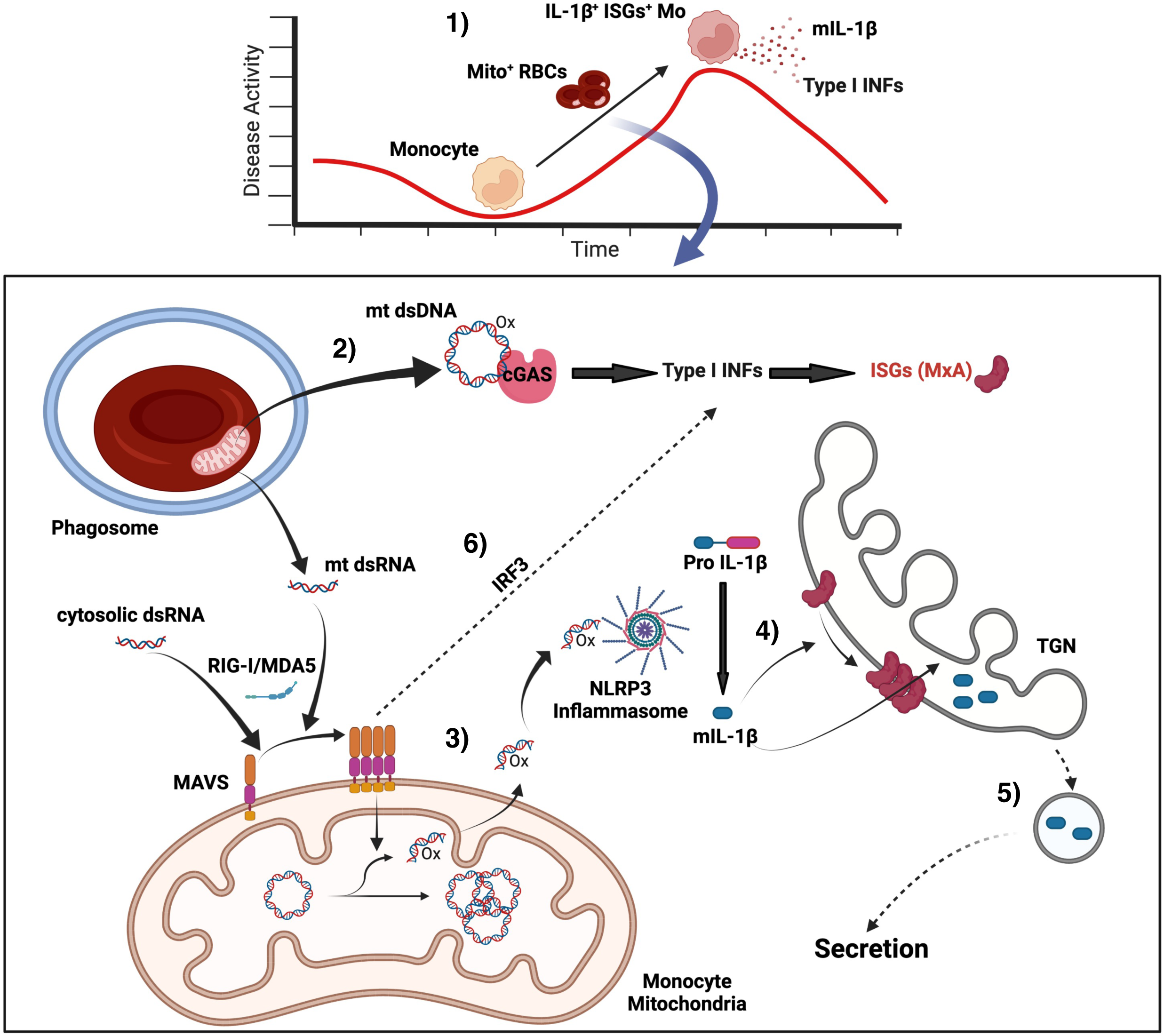
(1) In SLE patients with active disease, internalization of Mito^+^ RBCs induces Mo to co-express ISGs and mIL-1β. ISG expression depends on cGAS activation by Mito^+^ RBC-derived mtDNA (2), while mIL-1β production requires NLRP3 activation. Upstream of NLRP3, Mito^+^ RBC-derived mt dsRNA activates the RLRs/MAVS pathway, which induces the release of Mo-derived mtDNA fragments that bind NLRP3 (3). Importantly, mIL-1β triggers the oligomerization of MxA on the surface or the TGN (4), which favors the entry of this cytokine into a TGN-mediated unconventional secretory (vesicular?) pathway (5). (6) Cytosolic dsRNA of microbial origin (i.e. Poly I:C/Lyovec) could also activate these pathways via MAVS/IRF3.

**Table S1.**
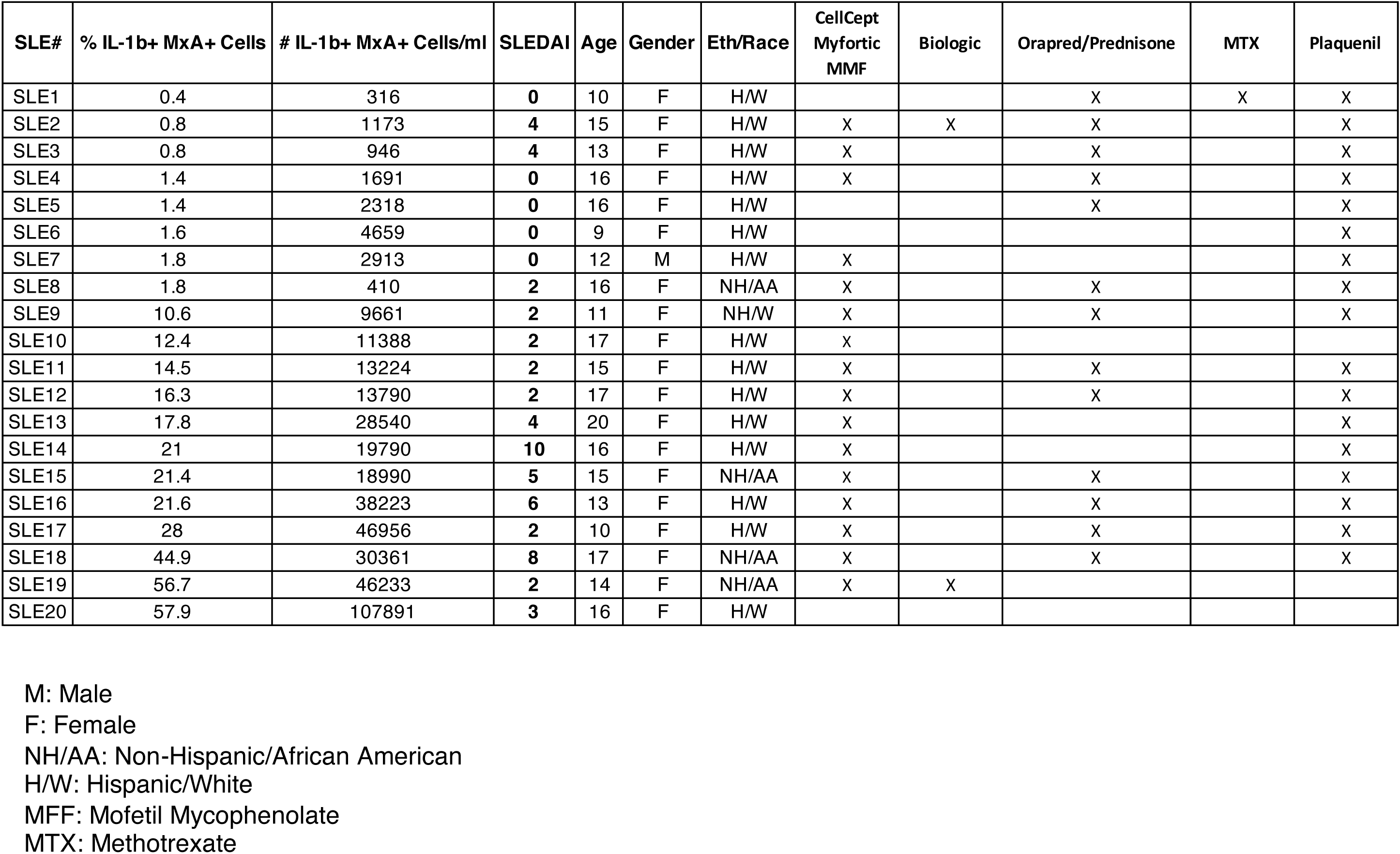

